# RANK is an independent biomarker of poor prognosis in estrogen receptor-negative breast cancer and a therapeutic target in patient-derived xenografts

**DOI:** 10.1101/2021.12.13.470911

**Authors:** Marina Ciscar, Eva M. Trinidad, Hector Perez-Montoyo, Mansour Alsaleem, Maria J. Jimenez-Santos, Michael Toss, Adrian Sanz-Moreno, Andrea Vethencourt, Gema Perez-Chacon, Anna Petit, Maria T. Soler-Monso, Jorge Gomez-Miragaya, Clara Gomez-Aleza, Maria Jimenez, Lacey E. Dobrolecki, Michael T. Lewis, Alejandra Bruna, Silvana Mouron, Miguel Quintela-Fandino, Fatima Al-Shahrour, Antonio Martinez-Aranda, Angels Sierra, Andrew R. Green, Emad Rakha, Eva Gonzalez-Suarez

## Abstract

Despite strong preclinical data, the therapeutic benefit of the RANKL inhibitor denosumab in BC patients, beyond its bone-related effects, is unclear. Here, we investigated the prognostic value of RANK expression and its functionality in human BC. We analyzed RANK and RANKL expression in more than 1500 BC cases (777 being estrogen receptor-negative (ER^-^)) from four independent cohorts. We confirmed that RANK is more frequently expressed in ER^-^ tumors, but it is also found in a subset of ER^+^ tumors. In ER^-^ BC, RANK expression was independently associated with poor outcome, especially in postmenopausal patients and those who received adjuvant chemotherapy. Gene expression analyses unraveled distinct biology associated with RANK in relation to ER expression and menopause, and evidenced enhanced RANK activation in ER^-^ postmenopausal tumors, together with regulation of metabolic pathways. Functional studies and transcriptomic analyses in ER^-^ RANK^+^ patients-derived orthoxenografts demonstrated that activation of RANK signaling pathway promotes tumor cell proliferation and stemness, and regulates multiple biological processes including tumor immune surveillance and metabolism. Our results demonstrate that RANK expression is an independent poor prognosis biomarker in postmenopausal ER^-^ BC patients and support the rational of using RANK pathway inhibitors in combination with chemotherapy in ER^-^ BC.

## Introduction

Breast cancer (BC) accounts for 10% of all types of cancers and 6.5% of global cancer-related mortality (1). BCs show a high pathological and biological heterogeneity, with considerable differences in histology, genetics and sensitivity to targeted therapies and chemotherapies. The histological expression of estrogen receptor (ER), progesterone receptor (PR), human epidermal growth factor receptor 2 (HER2) and Ki67 are determinant for BC prognosis and treatment (2, 3). Tumors expressing hormone receptors -luminal tumors-generally bear better prognosis than the rest of subtypes, whereas tumors lacking ER, PR and HER2 expression -the triple negative BC (TNBC)- have the worst outcome among BC subtypes and currently lack targeted therapies (4, 5). Despite recent advances in treatment (6), BC accounts for the main cause of mortality by cancer in women, highlighting the need of new prognosis markers and more optimized and individualized treatment approaches.

RANKL (*TNFSF11*) and its receptor RANK (*TNFRSF11A*) are candidates to explore as potential predictor biomarkers in BC. RANK is expressed on tumor cells in 40% of hormone receptor-negative tumors and 20% of the luminal tumors (7) and its expression is associated with a higher risk of relapse and death, given its abundance in TNBC (8). By contrast, RANKL is rarely found in tumor cells, being mostly restricted to a luminal A-like subset (8, 9), but has been found in the stroma surrounding the tumor (10).

Functionally, RANK signaling regulates mammary gland development and tumor initiation (10, 11). It mediates the proliferative response to progesterone in the mammary epithelia (12, 13) and the expansion of mammary stem cells and progenitors (10,14,15). RANK loss or RANKL inhibition blocks the occurrence of progesterone-driven tumors and attenuates tumorigenesis in genetically modified mouse models (10,15–17). Furthermore, the inhibition of RANK pathway induces tumor cell apoptosis and differentiation, reducing recurrence and metastasis in mouse mammary adenocarcinomas expressing RANK (17), supporting RANK signaling as a therapeutic target in BC.

RANK-RANKL involvement in bone remodeling prompted the development of denosumab, a fully human monoclonal antibody targeting and neutralizing RANKL. Denosumab is currently used for the treatment of patients with osteoporosis as well as for the prevention of skeletal-related events arising from bone metastases (18). In BC, adjuvant denosumab improved disease-free survival (DFS) in postmenopausal women with hormone receptor-positive BC (ABCSG-18 trial/NCT00556374) (19). In contrast, in the D-CARE study (NCT01077154), no survival differences associated with denosumab were found, neither in post nor in premenopausal women with high-risk early BC (20). These conflicting results highlight the need of further knowledge in the understanding of RANK signaling in BC and its therapeutic potential.

In this study, we have evaluated the potential value of RANK and RANKL as clinical predictors of prognosis in patients with different BC subtypes, and the functionality of RANK signaling in human BC.

## Results

### RANK expression in tumor cells associates with ER^-^ tumors and predicts poor survival in postmenopausal patients

To evaluate the potential of RANK and RANKL as prognosis biomarkers in BC, we analyzed the expression of both proteins in two independent tissue-microarray (TMA) collections containing all BC subtypes: *IDIBELL* (*IDB*) collection (n=404) (Martínez-Aranda et al. 2015) and *Nottingham primary series* (*NPS*) (n=1895 samples) (Green et al. 2013); a subset of *NPS* samples (n=298) were included in the Molecular Taxonomy of Breast Cancer International Consortium (*METABRIC*) (21).

RANK protein expression was observed in the tumor compartment in 18.3% of the samples in the *IDB* collection and 5.7% of the *NPS* (excluding *METABRIC* samples). RANK expression was detected in the stroma of approximately half of the cases, 55% and 46.3% from *IDB* and *NPS* cohorts, respectively (Fig. 1a-b). Tumor expression of transmembrane RANKL (tmRANKL) was found in only 4.6% (*IDB)* and 3.5% (*NPS)* of adenocarcinomas, in agreement with previous observations (10),(8), and was rarely found in the stroma (< 3%) (Fig. 1a-b). Fig. S1a shows the H-Score (H) for samples expressing RANK or tmRANKL in the tumor compartment.

**Fig. 1.**
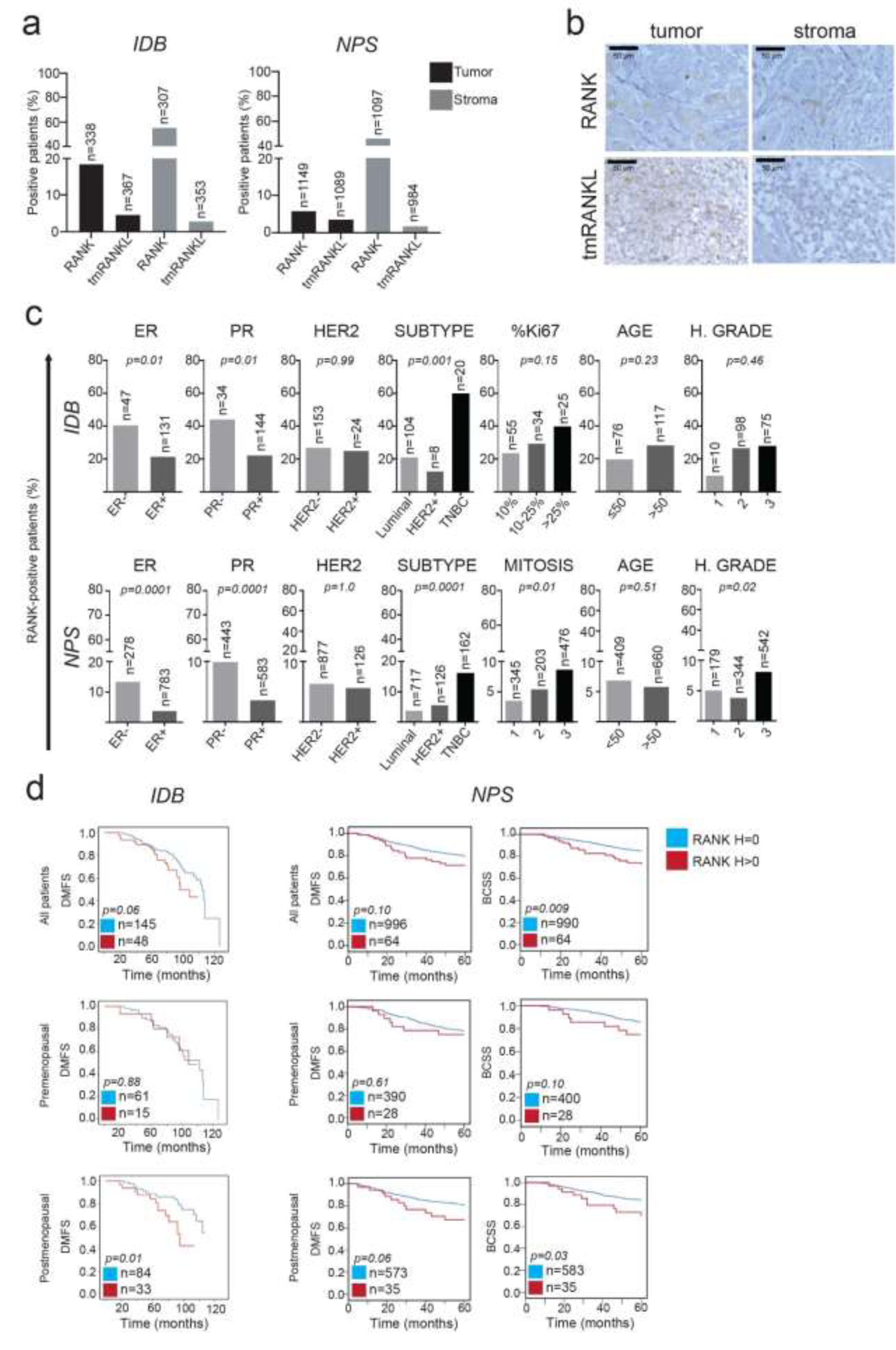
RANK is expressed in tumor and stromal cells of human BC and its expression in tumor cells associates with poor survival in postmenopausal patients. **(a)** Percentage of patients expressing tumor and stromal RANK or tmRANKL (H > 0) in BC samples from *IDB* and *NPS* collections. The total number of patients scored for RANK and tmRANKL expression is indicated. **(b)** Representative images showing RANK and tmRANKL protein expression in tumor and stromal cells in human BC determined by IHC. **(c)** Percentage of BC patients with RANK^+^ tumor according to the indicated clinicopathologic parameters in the *IDB* and *NPS* cohorts. The total number of patients analyzed per parameter and *p-values* (calculated using the Pearsońs ChiSquare test (Exact Sig. 2-Side)) are indicated. **(d)** DMFS and BCSS according to RANK expression (RANK^-^ (H = 0) or RANK^+^ (H > 0)) in all patients of the *IDB* and *NPS* collections and classified by menopause. The total number of patients analyzed per parameter and *p-values* (calculated using the Log-rank test (Mantel-Cox)) are indicated.

In both cohorts, RANK expression was significantly associated with ER/PR negativity and TNBC subtype, but not HER2, age, tumor size or stage. Furthermore, in the *NPS* collection, RANK expression was also associated with a higher mitosis rate and grade (Fig. 1c; Table S1). The low frequency of tmRANKL hinder associations with clinicopathologic parameters (RANKL expression associated with younger patients and low histological grade only in *NPS* collection) (Fig. S1b; Table S1). Similar expression patterns for RANK and RANKL were found in the *METABRIC* collection (Fig. S1c-e).

Patients with RANK^+^ tumors (H > 0) from *IDB* and *NPS* tended to have a poorer distant metastasis-free survival (DMFS) compared to those with RANK^-^ tumors (H = 0) (Fig. 1d, Table S1). Indeed, *NPS* patients with RANK^+^ tumors showed a worse 5-year BC-specific survival (BCSS) (Fig. 1d). Interestingly, RANK positivity associates with poor survival (DMFS and BCSS) in postmenopausal, but not in premenopausal patients from both collections (Fig. 1d, Table S1). In univariate survival analysis, patients in the *NPS* with RANK expression had a shorter 5-year BCSS compared to patients with no RANK expression and this association was maintained in multivariate Cox regression analyses when ER, tumor grade, stage and size were considered (Table S1). In the postmenopausal, but not in the premenopausal women, RANK expression associated with worse BCSS in univariate analyses, and worse BCSS and DMFS in multivariate analyses (Table S1). Tumor grade and stage, but not ER or tumor size, reached significance for all survival outcomes (Table S1). Altogether, our results confirm that RANK expression associates with ER^-^/PR^-^ tumors and TNBC subtype, and demonstrate that RANK expression can act as an independent biomarker of poor prognosis in postmenopausal BC patients.

### RANK does not associate with survival in ER^+^ BC, and tmRANKL expression associates with premenopausal patients

Given the different biology of ER^+^ and ER^-^ BC, we next aimed to address the clinicopathologic significance of RANK in each subgroup. In the ER*^+^* subsets, RANK in tumor cells was detected in 21.3% (*IDB)* and 3.7% (*NPS)* of samples, whereas tmRANKL only in 5.5% (*IDB)* and 3.9% (*NPS)*. RANK (73.2% in *IDB* and 51.1% in *NPS*), but not tmRANKL (2% in *IDB* and *NPS)*, was frequently found in the stroma (Fig. 2a). Tumor RANK expression in the ER^+^ subset of the *NPS* did not associate with any of the clinicopathologic factors or survival parameter analyzed, in neither premenopausal nor postmenopausal patients (Fig. 2b; Table S1). Tumor tmRANKL expression was associated with young women in the ER^+^ subset of *NPS* (Fig. S2a) and its expression correlated with better DFS (Fig. S2b).

**Fig. 2.**
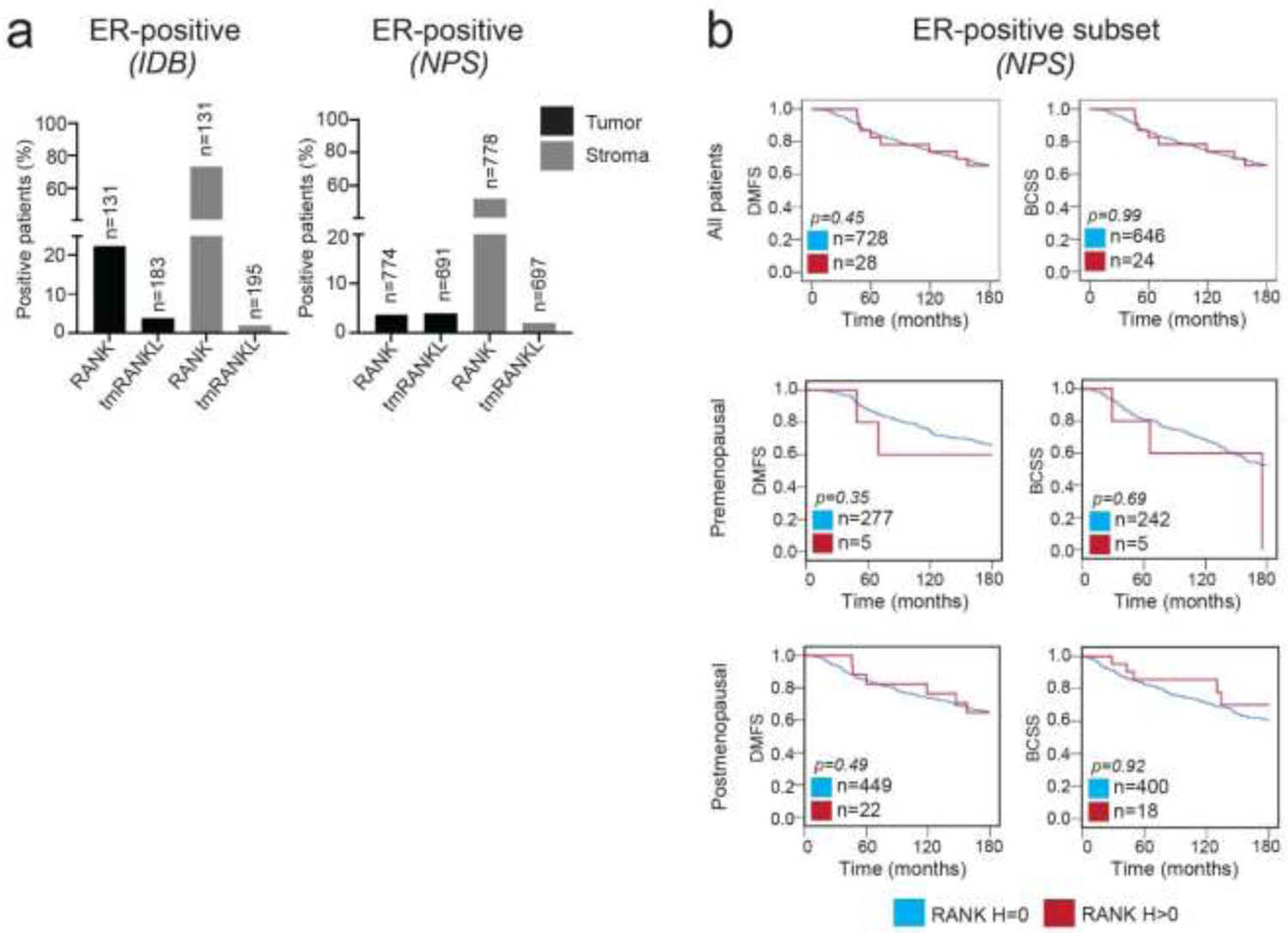
RANK tumor expression is not associated with survival in ER^+^ BC. **(a)** Percentage of tumor and stromal RANK or tmRANKL (H > 0) in the BC ER^+^ subset from the *IDB* and *NPS* collections. The total number of patients scored for RANK and tmRANKL proteins is indicated. **(b)** BCSS and DMFS in the ER^+^ subset from the *NPS* according to RANK expression, in all patients, premenopausal and postmenopausal (15 years of follow-up). The total number of patients analyzed per parameter and *p-values* (calculated using the Log-rank test (Mantel-Cox)) are indicated.

### RANK expression in ER^-^ tumors associates with poor response to chemotherapy and poor survival in postmenopausal patients

In the ER^-^ subsets, RANK was expressed in the tumor compartment in 40.4% and 13% of samples in *IDB* and *NPS* collections, respectively, and frequently found in the stroma (69.2% (*IDB*) and 43.7% (*NPS)*) (Fig. 3a). Again, tmRANKL was rarely found in the tumor or stroma of ER^-^ tumor samples (Fig. 3a). These RANK and RANKL expression patterns were confirmed in two additional and more recent collections of ER^-^ tumors: In the *ER-NEGATIVE ONLY* collection (396 ER^-^ tumors), tumor RANK and tmRANKL expression was found in 34% and 0.33% of samples, respectively (Fig. 3a); In the *TNBC* collection (n=66), we found that 30.3% of tumor samples were positive for RANK and 3.38% for tmRANKL; in the stroma RANK (65.2%), but not tmRANKL (6.7%), was commonly expressed (Fig. 3a).

**Fig. 3.**
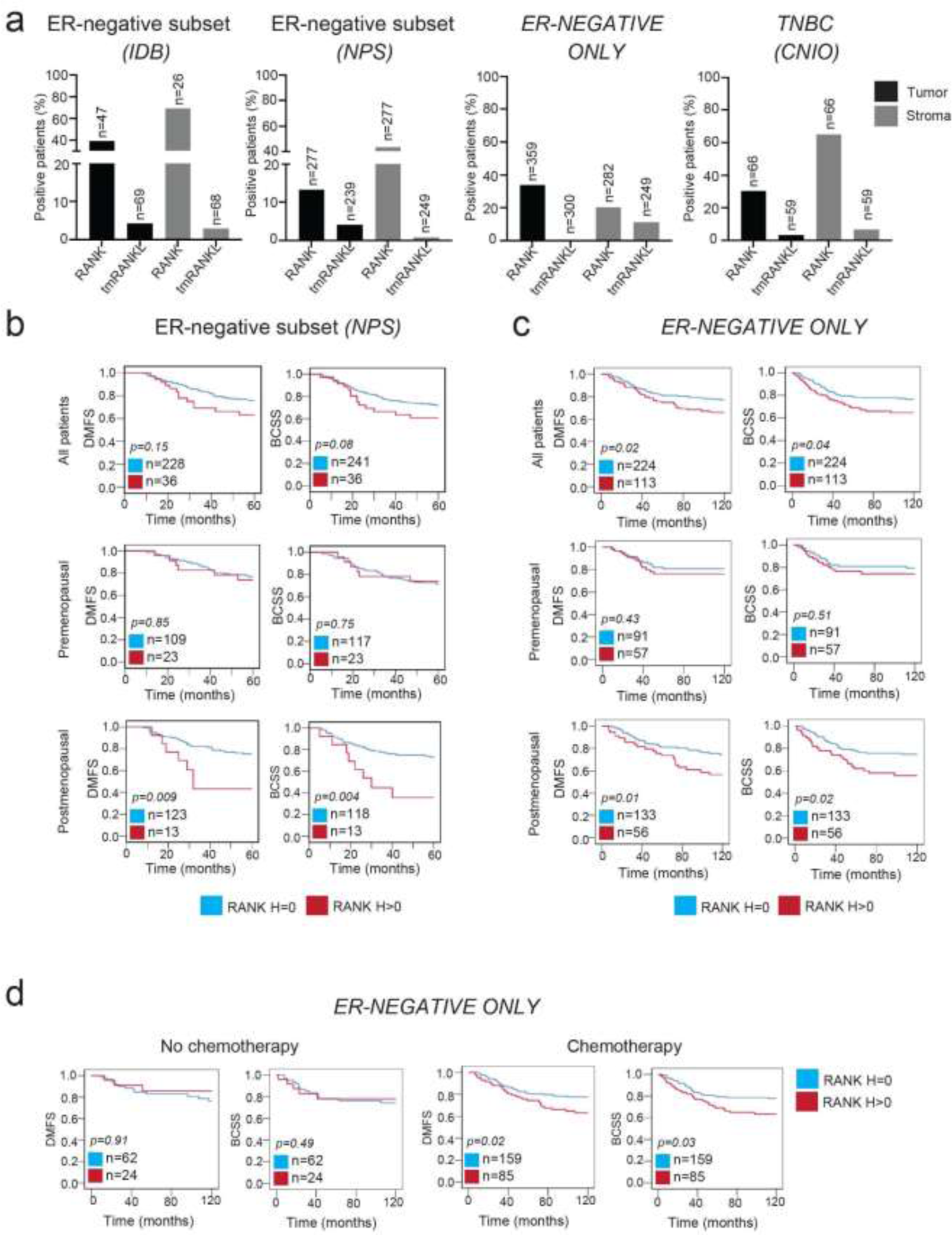
RANK tumor expression is associated with a reduced survival in postmenopausal patients with ER^-^ tumors and poor response to chemotherapy. **(a)** Percentage of patients expressing tumor and stromal RANK or tmRANKL (H > 0) in the ER^-^ subset from the *IDB* and *NPS* collections, and in two additional cohorts containing exclusively ER^-^ samples: *ER-NEGATIVE ONLY* (Nottingham) and *TNBC* (CNIO). The total number of patients scored for RANK and tmRANKL protein expression is indicated. **(b)** DMFS and BCSS according to RANK expression in the ER^-^ subset from NPS (5 years of follow-up) in all patients and in premenopausal and postmenopausal patients. **(c)** DMFS and BCSS according to RANK expression in all patients, premenopausal and postmenopausal patients with ER^-^ tumors in *ER-NEGATIVE ONLY* collection (10 years of follow-up). **(d)** DMFS and BCSS in the *ER-NEGATIVE ONLY* collection after chemotherapy or not, according to RANK expression (10 years of follow-up). **(b, c, d)** The total number of patients analyzed per parameter and *p-values* (calculated using the Log-rank test (Mantel-Cox)) are indicated.

Tumor RANK expression in ER^-^ tumors was not associated with any of the clinicopathologic factors analyzed (Fig. S3a, Table S1). RANK expression tended to be associated with worse DMFS and BCSS in the *NPS* ER^-^ subset, although differences did not reach significance (Fig. 3b). However, when menopausal status was considered, RANK expression was associated with poorer survival at 5 years in the ER^-^ subset from postmenopausal, but not premenopausal, patients; the association of RANK with poor prognosis in this cohort was observed up to 15/20 years (Fig. 3b, Table S1). As frequency of RANK positivity in this collection was low (13%), we confirmed this finding in the *ER-NEGATIVE ONLY* collection, where RANK detection was higher (34%) (Fig. 3a). Importantly, in the *ER-NEGATIVE ONLY* collection, patients with RANK^+^ tumors also showed a significant poorer 10-year survival compared to patients with RANK^-^ tumors. RANK association with worse survival was observed only in postmenopausal women (Fig. 3c, Table S1). Cox regression analyses demonstrated that RANK expression was an independent factor of worse 10-years DMFS and DFS in all ER^-^ patients, as well as in postmenopausal, but not in premenopausal, patients in the *ER-NEGATIVE ONLY* collection. Tumor stage was independently associated with the three survival parameters analyzed (Table S1).

Moreover, patients with ER^-^ RANK-expressing tumors showed poorer survival after adjuvant chemotherapy than those lacking RANK, while no survival differences associated with RANK were found in the absence of chemotherapy (Fig. 3d, Table S1). Similarly, in the *CNIO TNBC* collection (Fig. S3b) tumors expressing RANK tended to have worse survival in patients receiving chemotherapy, particularly to regimens containing taxanes (Fig. S3b, Table S1). The low frequency of tumor tmRANKL positivity in the ER^-^ subsets prevented reliable associations with any parameter (Fig. S3c-d). Taken together, these results point out the importance of RANK expression in ER^-^ tumors as a biomarker of poor prognosis, mainly in postmenopausal ER^-^ BC.

### Distinct RANK biology according to ER expression and menopause

Our previous results demonstrate that RANK expression in tumor cells is a biomarker of poor prognosis in ER^-^ but not in ER^+^ tumors, and in postmenopausal, but not in premenopausal patients, suggesting putative differences in RANK tumor biology in both tumor subtypes. Thanks to the availability of gene expression data from the *METABRIC* dataset, we analyzed pathways differentially regulated between RANK^+^ and RANK^-^ tumors in ER^+^ and ER^-^ BC and in pre and postmenopausal patients.

Results from gene set enrichment analyses (GSEA) revealed 67 pathways associated with RANK protein expression in ER^+^ tumors and 17 in ER^-^ tumors (FDR < 0.25), with no overlap between them. Several pathways related to metabolism (Normalized Enrichment Scores (NES) < 0) and immunity (NES > 0) associated with RANK in ER^-^ tumors (Fig. S4 and Table S2). In ER^+^ tumors, several pathways related with DNA replication and gene transcription were negatively associated with RANK expression (NES < 0) (Fig. S4a and Table S2). No common pathways associated with RANK expression (FDR < 0.25) were observed between tumors from premenopausal and postmenopausal patients. Importantly, RANK expression in postmenopausal tumors positively associated (NES > 0) with multiple pathways related to TNF/NFκB signaling, including RANKL pathway itself (Fig. S4 and Table S2); suggestive of a more active RANK signaling in the breast tumors from postmenopausal, than in those of premenopausal patients, as happens in the bone (22). In ER^-^ premenopausal, RANK only associated with two pathways, FGFR2 and hedgehog signaling, meanwhile in ER^-^ postmenopausal RANK expression positively associated with 21 pathways, including positive associations with several TNF/NFκB signaling pathways and immune pathways, and negative associations with multiple metabolic pathways, insulin/IGF1 signaling, fatty acid metabolism and mTOR (Fig. S4 and Table S2). Together, these results highlight the different biology of RANK signaling according to ER status and menopause and suggest that enhanced RANK signaling in postmenopausal tumors and regulation of tumor cell metabolism may contribute to the association of RANK expression with poor prognosis in ER^-^ postmenopausal tumors.

### RANK is expressed and functional in ER^-^ BC patient-derived orthoxenografts (PDXs)

Our results support the relevance of RANK expression as a prognosis factor in human BC, particularly in ER^-^ disease and response to chemotherapy. Despite encouraging results in mouse models (17), direct demonstration of the functionality of RANK signaling in human BC is lacking. To this end, we analyzed human *RANK* and *RANKL* gene expression in several collections of BC PDXs (23–28) derived from human BC (Fig. 4a). *RANK* mRNA was detected in all models tested, with expression levels varying 10-fold between different models. Tumors with the highest levels of *RANK* mRNA expression were found in the ER^-^ tumors, in accordance with previous findings (29). Meanwhile, *RANKL* gene expression was low or even undetectable in most PDX models, with some exceptions (Fig. 4a). RANK protein expression was found in 40% of ER^-^ and 14.3% of ER^+^ from the 76 PDX analyzed, recapitulating the clinical patterns (Fig. 4b, Table S3). We detected RANK protein expression in some of the selected PDXs (#) with high or intermediate *RANK* mRNA levels, whereas tmRANKL protein was only detected in HCI-001 and STG139-M (Fig. 4c). Next, we analyzed NFκB activation upon RANKL stimulation *in vitro,* as it is the main pathway regulated by RANK in BC, and we found it associated with RANK^+^ tumors (Fig. S4). A clear enhanced phosphorylation of IkBα and/or p65 after RANKL treatment was observed only in the models AB521-X, BCM-3277 and STG139-M ( Fig. 4d, Fig. S5a). Gene expression analyses of several RANK/NFκB targets confirmed RANK pathway activation in BCM-3277, AB521-X and STG139-M (Fig. S5b). Although BCM-3277 derived from a human luminal tumor, ER expression was not detected in the PDX, and PAM50 analyses classified it as “basal-like”. Analyses of surface markers, CD44, CD24, EpCAM, CD133 and CD10, confirmed similar expression patterns to those reported in BC PDXs (Fig. S5c) (27).

**Fig. 4.**
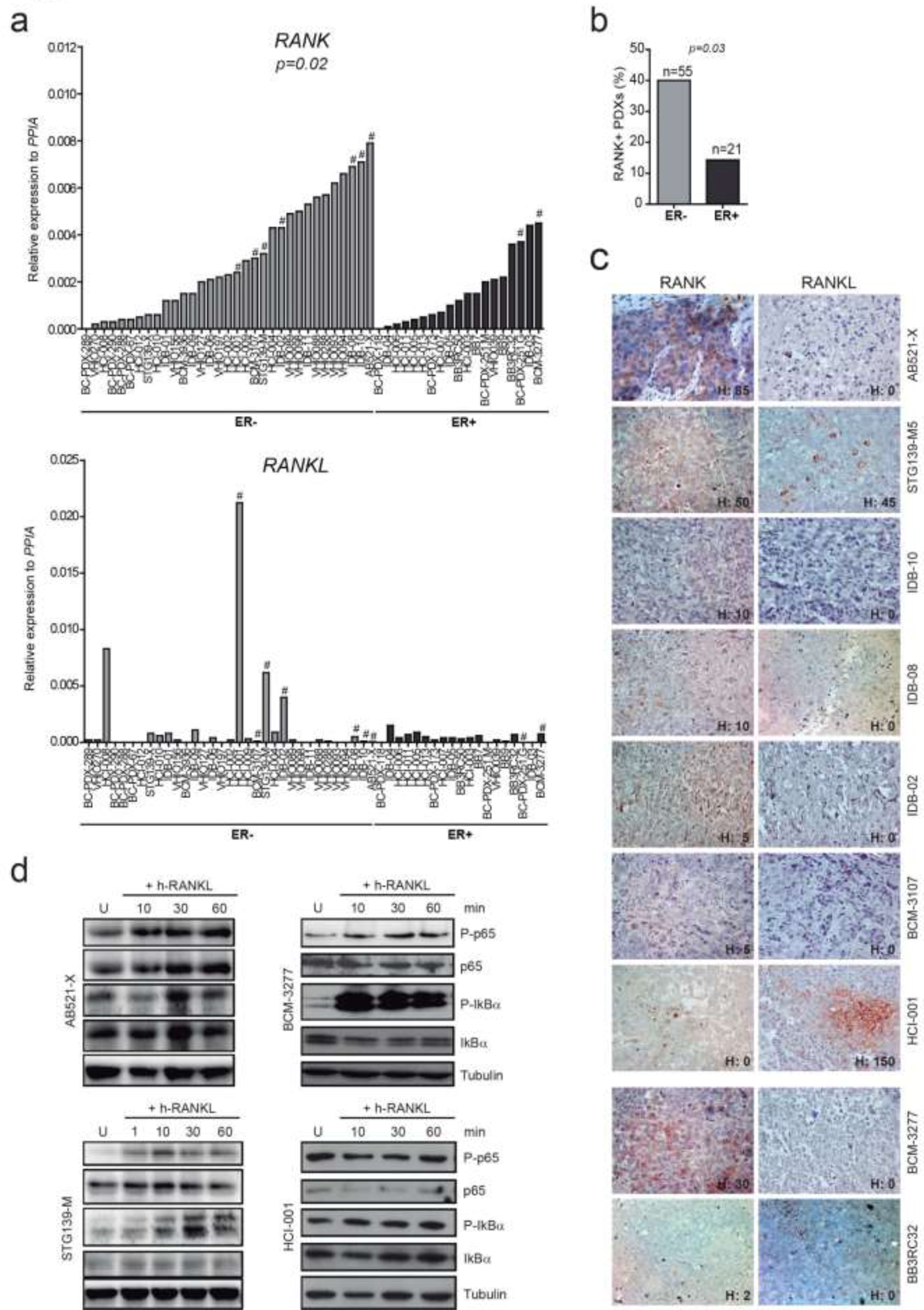
RANK is expressed and functional in BC PDXs. **(a)** RANK and RANKL mRNA expression levels relative to PPIA in the indicated BC PDXs, organized according to ER status in the human tumor of origin and RANK mRNA expression. Two tailed t-student test was used to evaluate the RANK/RANKL differential expression between ER^-^ and ER^+^ BC PDXs. # Indicates models where RANK and RANKL expression were analyzed by IHC. **(b)** Percentage of PDXs expressing RANK protein according to ER expression. Total number of independent PDXs analyzed is shown. *P-value* was calculated using a two tailed t-student test. **(c)** Representative images of RANK and RANKL protein expression in BC PDXs detected by IHC. H-Score (H) of the models (and not of the picture) are shown. A total of 3-5 independent tumors per PDX, were scored for RANK. Pictures are ordered according to RANK mRNA expression levels and subtype in the human samples of origin. **(d)** Western blot analyses of P-p65, P-IKBα and corresponding total proteins after the minutes (min) of RANKL stimulation in the indicated PDXs. Tubulin was used as a loading control.

### RANK pathway promotes tumor cell proliferation and stemness in ER^-^ BC PDXs

Next, we evaluated the functional consequences of RANK pathway modulation *in vivo* in three independent BC PDX models responsive to RANKL. Tumor-bearing NOD SCID Gamma (NSG) mice were randomized for treatment with h-RANKL, the inhibitor RANK-Fc, denosumab (in the STG139-M model as it expresses hRANKL) or mock treatment (controls) for 4 weeks (Fig. S6a). Mice treated with RANKL showed increased levels of the bone remodeling marker,Trap5b, while those treated with RANK-Fc, which binds mouse and human RANKL had lower Trap5b (Fig. S6b). RANKL inhibition slightly attenuated tumor growth, while RANKL stimulation modestly increased tumor growth in AB521-X (Fig. 5a).

**Fig. 5.**
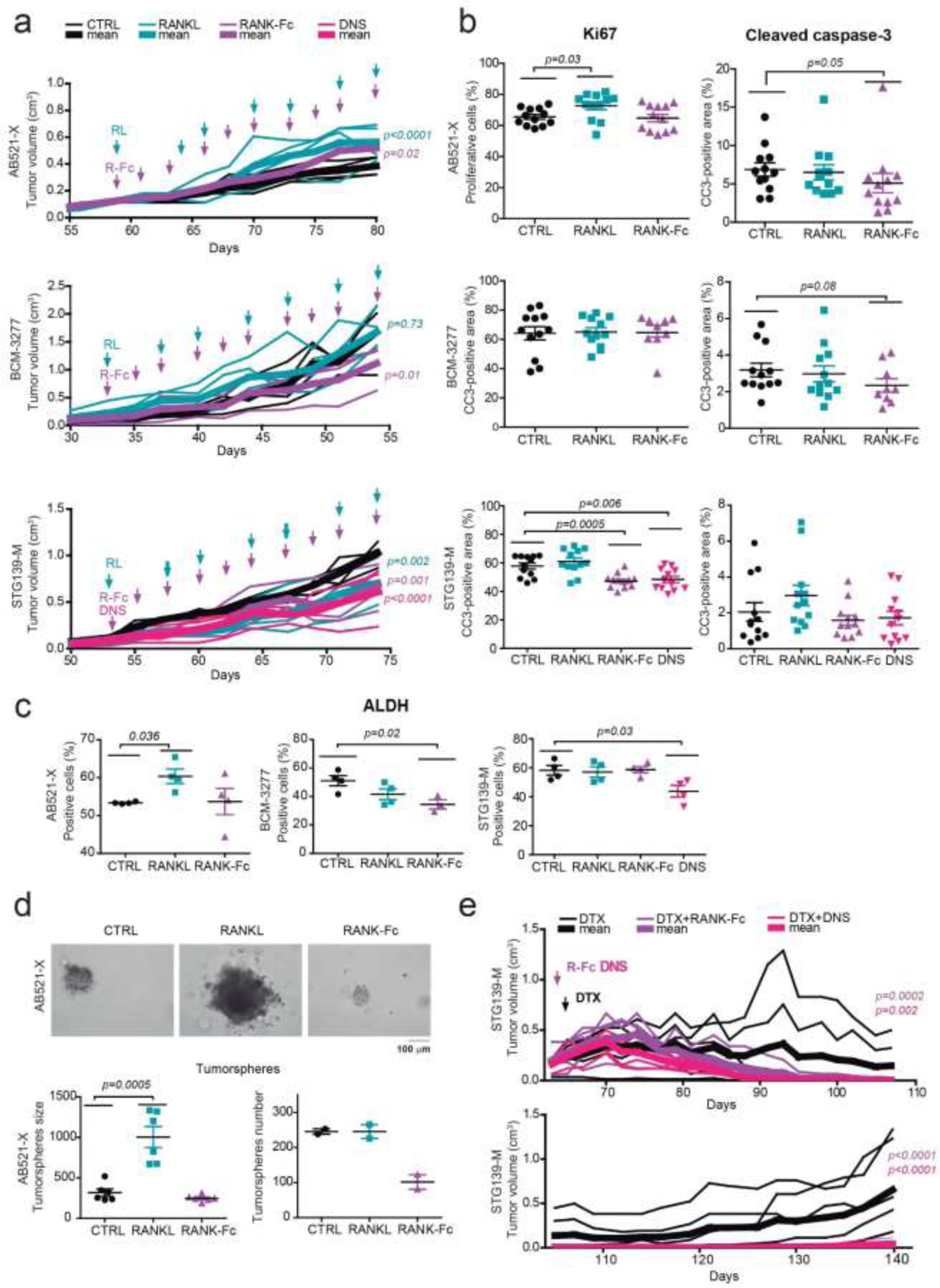
RANK signaling regulates tumor cell proliferation, stemness and chemotherapy response in BC PDXs. **(a)** Tumor growth curves ((π x length x width2)/6) of the indicated PDXs after treatment with RANKL, RANK-Fc, denosumab (DNS) or mock treatments (CTRL). Treatment started as indicated by the arrows following the scheme in Fig. S6a. Each thin curve represents one single tumor, and each thick curve represents the mean of all tumors implanted. Linear regression analysis was performed and two-tailed *p-value* is shown. **(b)** Percentage of cells positive for Ki67 and the cleaved caspase-3 positive area in tumors of the indicated PDXs collected 24 hours after last treatment (Fig. S6a). Each dot represents one picture. Three representative pictures per tumor were quantified and at least 3-4 tumors per condition were analyzed. Two-tailed t-test *p-values* are shown. **(c)** Percentage of cells with ALDH+ activity in tumors isolated from the indicated PDXs, collected 24 hours after last treatment (Fig. S5a). Each dot represents one tumor. The treated tumors were compared with controls used a two-tailed t-student test. **(d)** Representative images of secondary tumorspheres derived from tumor cells from in vivo treatments. Total number of secondary tumorspheres (each dot represents a tumor) and tumorsphere size (each dot represents an average by tumor) are shown. *P-value* was calculated using a two tailed t-student test **(e)** Tumor growth curves ((π x length x width 2)/6) of indicated PDXs after treatment with docetaxel (DTX, 20 mg/kg, once per week) in combination with RANK-Fc or denosumab (DNS). Treatment started when indicated by the arrows. Each thin curve represents one single tumor and each thick curve represents the mean of all tumors implanted. Linear regression analysis was performed and two-tailed *p-value* is shown.

RANKL treatment increased tumor cell proliferation (ki67) in the model AB521-X, which had the highest RANK protein expression. Conversely, inhibition of RANKL by RANK-Fc or denosumab decreased tumor cell proliferation in the STG139-M model and not in the BCM-3277 model (Fig. 5b, Fig. S6c). A slight decrease in tumor cell apoptosis (cleaved-caspase 3) after RANKL inhibition was observed in AB521-X (Fig. 5b, Fig. S6c).

An increase in aldehyde dehydrogenase (ALDH) activity after RANKL treatment was observed in the AB521-X model and RANKL inhibition reduced ALDH activity in the other two models (Fig. 5c), in accordance with RANK signaling enhancing BC stemness (17). RANKL treatment led to an increase in tumorsphere size in AB521-X model, whereas its inhibition reduced the number of secondary tumorspheres in the AB521-X and the BCM-3277 models (Fig. 5d, Fig. S6d), supporting a decrease in BC stemness. STG139-M tumor cells did not grow as tumorspheres when plated in suspension. Together, these results demonstrate the functionality of RANK signaling in human BC and suggest that inhibition of RANK signaling in human BC can reduce tumor cell proliferation and stemness.

### RANKL inhibitors improve the response to docetaxel in ER^-^ BC PDXs

Clinical analyses (Fig. 3d and Fig. S3b) evidenced that RANK expression in ER^-^ tumors was associated with poor survival after chemotherapy. To directly test whether RANK pathway inhibitors could improve response to chemotherapy, tumor-bearing NSG mice were randomized for treatment with docetaxel alone or in combination with RANK-Fc or denosumab (for STG139-M) (30): docetaxel treatment was interrupted when tumor diameter decreased below 3 mm. The three PDX models were sensitive to docetaxel, but increased benefit was observed when adding RANKL inhibitors (Fig. 5e and Fig. S6e). In the STG139-M model, docetaxel treatment could not be interrupted in some tumors, and they rapidly regrew even in the presence of docetaxel. In contrast, when denosumab or RANK-Fc were added to docetaxel, all tumors disappeared, and no tumor relapses were observed even 60 days after interruption of docetaxel treatment (Fig. 5e). Together, these results demonstrate that inhibition of RANK signaling can reduce tumor cell proliferation and stemness and improve response to chemotherapy in ER^-^ BC patients.

### RANK signaling in BC PDXs regulates pathways involved in cell proliferation, metabolism, stemness and immunity

Considering BC heterogeneity, it is important to identify the molecular mechanisms underlying the response to denosumab in human BC cells. To this aim, RNAseq analysis was performed after modulation of RANK signaling in the three PDX models (Table S4). GSEA results demonstrated the strong impact that modulation of RANK signaling caused in these BC PDX with approximately 200 pathways differentially regulated (FDR < 0.25) in each PDX after RANKL, RANK-Fc or denosumab treatment (Table S4). Despite the great heterogeneity in BC and between the PDX models, most pathways modulated by RANKL and RANK-Fc were shared between the three different PDX models (FDR < 0.25) (Table S4). The top-ranked RANKL-driven pathways shared by the three models (NES > 0) were related to TNF/NFκB signaling, immunity (IL2/STAT5, complement, interferon gamma response), proliferation (G2M checkpoint, DNA repair) as well as cell adhesion and stemness (WNT signaling). Metabolic pathways (i.e. insulin signaling, reactive oxygen species, glycolysis, adipogenesis) were positively associated (NES > 0) with RANK-Fc treatments (Fig. 6 and Table S4). Pathways related to adhesion, immunity and estrogen response were found in both RANKL- and RANK-Fc-treated tumors. As expected, RANK-Fc and denosumab modulated the same pathways in STG139-M (Fig. 7a and Table S4). Next, we compared pathways regulated in PDX after pharmacological treatments with RANKL/RANK-Fc, with those associated with RANK in BC clinical samples (Fig. S4 and Table S2). Importantly, there was a stronger overlap with ER^-^ BC and even more with ER^-^ postmenopausal, reinforcing that RANK signaling is more active in ER^-^ postmenopausal patients, where is a key regulator of tumor cell immunity and metabolism.

**Fig. 6.**
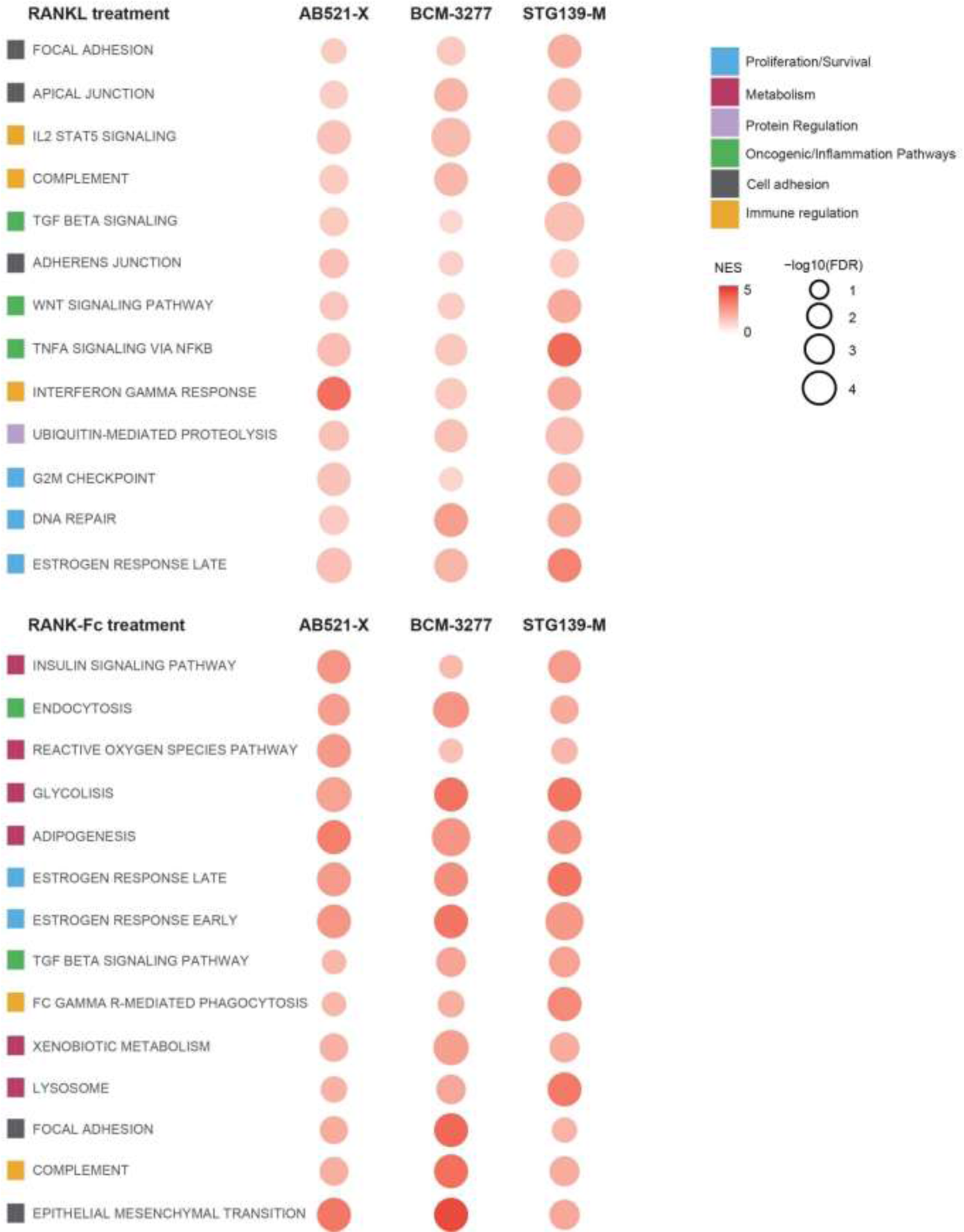
RANK signaling regulates proliferation, stemness, adhesion, metabolism and immunity in BC PDXs. The bubble matrix represents GSEA results of associated genes after in vivo treatment with RANKL and RANK-Fc in NSG mice, which are common for the 3 PDX models studied. The matrix illustrates NES and *FDR* values. The color scale represents the NES: the more intense the color, the more positive it is. In addition, the size of the bubbles is proportional to the -log10 of the FDR. Thus, the bigger the dot, the smaller the FDR. For those signatures with an *FDR = 0* after 1000 permutations, we assigned an *FDR =* 10*^-3* for visualization purposes. The signatures selected for this plot belong to Hallmark, Biocarta, Reactome and KEGG collections and have a reported *FDR < 0.05* and a NES > 0 for all PDX models. The color legend indicates the main biological process associated to each signature.

To confirm the relevance of gene expression changes observed in PDXs and those identified in BC patients, GSEA was performed with the genes modulated by denosumab in early BC from the D-BEYOND clinical trial (NCT01864798) (31). Importantly, the genes up-regulated by denosumab significantly associated with RANK signaling inhibition in the three PDX models (Fig. S7b, Table S4). Together, these results evidence the pleiotropic effects of RANK signaling in human BC tumor cells and suggest that denosumab will impact, not only tumor cell proliferation and stemness, but also immunity, cell adhesion and metabolism.

### Experimental procedures

#### Tissue Microarray (TMA) staining and scoring

RANK and tmRANKL expression were evaluated in TMAs from five different cohorts of BC patients. *IDB* TMA (donated by A. Sierra (IDIBELL, Spain), contains 404 BC samples and clinicopathologic information from 314 patients (24-88 years old) diagnosed between 1989 and 2009. Follow-up ranged from 8 to 146 months (mean: 76.6 months). Metastasis relapse occurred in 43.4% (138/318) of patients; of these, 84 patients (60.9%) developed brain metastasis, 47 (34.1%) lung metastasis, 54 (39.1%) liver metastasis, 40 (29.0%) non-regional lymph node metastasis and 89 (64.5%) bone metastasis. Just over half (56.6%; 180/318) of the patients had no metastatic progression after a minimum follow-up of 5 years. *NPS* TMA is a well-characterized cohort of unselected early-stage (I-III) primary operable invasive BC from patients aged 70 years or younger, enrolled into the Nottingham Tenovus Primary Breast Carcinoma Series between 1990 and 1997 (*n* = 1895), and managed in accordance with a uniform protocol; a subset of cases (*n* = 298) were included in the *METABRIC* study (21), where gene expression data is available. Outcome data include survival status, survival time, cause of death, development, and time to locoregional recurrence and distant metastasis (DM). BCSS is defined as the time (in months) from the date of primary surgery to the date of breast cancer-related death. DMFS is defined as the time (in months) from the date of primary surgery to the appearance of DM. Treatments include chemotherapy (CMF) or endocrine therapy. At that time patients with HER2^+^ tumors had no access to trastuzumab. Positive ER status was defined as > 1% of tumor cells expressing ER. Positive HER2 status was defined using immunohistochemistry as HER2 3+. Histological grade was assessed based on the Nottingham Grading System (32, 33). Other clinicopathologic factors such as ER, PR and/or HER2 expression, proliferation rate (Ki67 expression or mitosis), vascular invasion, as well as patient age and survival analysis were analyzed before including the samples into the TMAs (34). Two additional collections of ER^-^ tumors were analyzed, the Nottingham *ER-NEGATIVE ONLY* cohort (1998-2006), which contains 396 samples, and the *CNIO TNBC*, a small collection of 66 patients with TNBC with 40-50% of relapse, generated by Dr M. Quintela-Fandino (CNIO, Spain).

RANK or tmRANKL staining was scored for intensity (on a scale of 0 to 3; 0 = no staining, 1 = weak, 2 = moderate, 3 = intense) and positive cell percentage (on a scale of 0 to 100%) within tumor cells or surrounding stroma for each TMA core sample. The sum of multiplying staining intensity by positive area is in the H-Score value, ranging from 0 to 300. TMA cores with less than 30% of the core area were discarded. Patients were stratified according to RANK or tmRANKL H-Scores as being protein-positive (H > 0) or protein-negative (H = 0). As TMA samples are enriched in tumor cells, the stroma content was not always present or representative and H-Score was not calculated. Total number of scorable samples for each of the collections and stainings is indicated in the corresponding figure.

### Generation of PDXs

IDB PDXs were generated by orthotopic transplantation of human fresh tumor tissue or injection of metastatic cancer cells isolated from pleural effusions into the cleared mammary fat pad of immunodeficient mice, as described (27). The rest of the PDXs were obtained through collaboration with Dr V. Serra and Dr J. Arribas (Vall d’Hebron Institute of Oncology), Dr A. Welm (Huntsman Cancer Institute), Dr M. T. Lewis (Baylor College of Medicine), Dr A. Bruna and Dr C. Caldas (Cancer Research UK Cambridge Institute) and Dr R. Clarke (Manchester Breast Centre).

PDXs were maintained by consecutive rounds of transplantation with tumor pieces. In the case of the B3277 model, mice were treated with 17β-estradiol at 8 µg/ml (Sigma) in drinking water. Mice were kept in individually ventilated and open cages and food and water were provided ad libitum. Cages, bedding, food and water were all autoclaved. Euthanasia was performed by CO_2_ inhalation.

### Tumor cell isolation

Single cells were isolated from tumors as described previously (35). Briefly, fresh tissues were mechanically dissected with a McIlwain tissue chopper and enzymatically digested with appropriate medium (DMEM F-12, 0.3% collagenase A, 2.5 U/ml dispase, 20 mM HEPES, and 100 U/ml penicillin/100 μg/ml streptomycin) 60 minutes at 37 °C. Samples were washed with Leibowitz L15 medium/10% FBS between each step. Erythrocytes were eliminated with hypotonic lysis buffer, and fibroblasts were excluded by incubation with DMEM F-12/10% FBS 1 hour at 37 °C. Single epithelial cells were isolated by treating with trypsin 2 minutes at 37 °C. The cell suspension was filtered with 40 mm filters and counted.

### RANKL, RANK-Fc, denosumab and docetaxel treatments in vivo

Dissociated tumor cells mixed 1:1 with Matrigel Basement Membrane (BD Biosciences) were transplanted orthotopically in the inguinal mammary gland of 10/12-week old NSG mice and when tumors reached 5 mm of diameter mice were randomized for mock, h-RANKL (0.75 mg/kg, 4-6 doses, twice per week; Amgen Inc), h-RANK-Fc treatment (10 mg/kg, three times per week; Amgen Inc), denosumab (10 mg/kg, three times per week; XGEVA ®). After 24 hours that the treatment was completed, mice were sacrificed and tumors were surgically extracted. Docetaxel (Hospira/Actavis, 20 mg/kg) was administered once per week together with dexamethasone (0.132 mg/kg, Merck), to reduce the inflammation caused for the chemotherapeutic. All the drugs were intraperitoneally injected. Tumor development was monitored once per week. Tumor volume was calculated as: π x length x width^2^/6 in cm. In combined docetaxel treatment mice were sacrificed once relapses reached 10 mm of diameter.

### Flow Cytometry

Single tumor cells were resuspended and incubated in blocking solution (PBS containing 2% FBS, 2 mM EDTA and IgG blocking reagent (Sigma)) for 10 minutes on ice. Then, the cells were labeled with fluorophore-conjugated antibodies against: CD24-PE (ML5, 555428), CD44-APC (G44-26, 555478), EpCAM-APC (EBA-1, 347200), CD10-PECy5 (HI10a, 555376) and CD49f-A647 (GoH3, 555735) from BD Pharmingen and CD133/1-PE (AC133, 130-098-826) from Miltenyi Biotec. Mouse cells were excluded in flow cytometry using H2Kd-PECy7 (SF1-1.1, 116622 from BioLegend). Gating was based on “Fluorescence Minus One” controls. Single tumor cells were assessed for their ALDH activity using the ALDEFLUOR™ Kit for ALDH Assays system (01700 from STEMCELL Technologies), following the manufacturer’s procedures. DAPI (Thermo Fisher Scientific) was added in the different antibody combinations to discriminate dead cells. A population of 10,000 living cells was captured in all flow cytometry experiments. Flow cytometry analysis was performed using a Gallios flow cytometer (Beckman Coulter). Data were analyzed using the FlowJo software.

### Enzyme-linked immunosorbent assay (ELISA)

TRAP 5b activity was measured in mouse serum according to the manufacturer’s instructions (IDS).

### Tissue histology and immunostaining

Three-micrometer sections were cut and immunohistochemistry of hRANK and hRANKL was performed as described (7, 10). RANK antigen retrieval was carried out with the Diva Decloaker buffer (Biocare Medical), 90 °C, 14-16 hours whereas sodium citrate buffer (0.01 M, pH=6) was used for RANKL. Protein block was done with TNB Blocking Buffer (PerkinElmer). Anti-human RANK monoclonal antibody (N-1H8; Amgen, 5 μg/mL) and anti-human RANKL monoclonal antibody (Amgen, M366, 1.85 μg/mL), anti-Ki67 (SP6, Abcam) and the -cleaved caspase-3 (Asp175, Cell Signaling) antibodies were used. VECTASTAIN® Elite® ABC-HRP Kit (Vector Laboratories) was used to amplify the RANK, RANKL and cleaved caspase-3 staining. Images were analyzed with Fiji software (36).

### Tumorsphere assays

Primary tumorspheres were derived by plating 100,000 tumor cells/mL in 2 mL of DMEM F-12, containing EGF 10 ng/ml, FGF 10 ng/ml, B-27™ Supplement (Gibco), 4 mg/mL heparin and penicillin/streptomycin in ultra-low attachment plates (Corning® Costar®). After 14 days, single tumors were isolated by 5 minutes treatment with PBS-EDTA 1 mM and 5 minutes of trypsin at 37 °C and plated for secondary tumorsphere formation at a concentration of 10,000 cells/mL in triplicates and counted 14-21 days later. Media was renewed weekly including RANKL (500 ng/ml; Amgen Inc) or RANK-Fc (1 µg/mL; Amgen Inc) as needed. Three different pictures were taken for each well and measured the diameter of the tumorspheres with Fiji software.

### RANKL stimulation in vitro

PDX single tumor cells were embedded in Corning™ Matrigel™ Growth Factor Reduced Basement Membrane Matrix (Corning™ 356238), plated in DMEM F-12 with B-27™ Supplement, EGF 10 ng/ml, hydrocortisone 0.5 μg/ml, insulin 5 μg/ml, cholera toxin 100 ng/ml, and penicillin/streptomycin. Cells were stimulated, or not, with h-RANKL (500 ng/mL; Amgen Inc) during 24 hours prior to gene expression analyses.

### Quantitative Reverse Transcription PCR (Q-RT-PCR)

Total RNA was isolated from tumor pieces using TRIzol (Thermo Fisher Scientific) or Maxwell® RSC simplyRNA Tissue Kit (AS1340 Promega). One μg of RNA was reverse-transcribed into cDNA using 200 U Superscript II plus random hexamer oligos (Invitrogen) and *RANK* and *RANKL* expression was analyzed relative to *PPIA* with LightCycler® 480 Probes Master (Roche, 04707494001) and a LightCycler® 480 thermocycler (Roche). The primers sequences used were: *PPIA-UPL* (Fw: ATGCTGGACCCAACACAAAT; Rv: TCTTTCACTTTGCCAAACACC), *TNFRSF11A-UPL* (Fw: GCAGGTGGCTTTGCAGAT; Rv: GCATTTAGAAGACATGTACTTTCCTG), *TNFSF11-UPL* (Fw: TGATTCATGTAGGAGAATTAAACAGG; Rv: GATGTGCTGTGATCCAACGA).

Gene expression in organoid cultures were evaluated using SYBR Green Master I (Roche, 04887352001), LightCycler® 480 thermocycler (Roche). The primers used were: *PPIA* (Fw: ATGGTCAACCCCACCGTT; Rv: TCTGCTGTCTTTGGGACCTTG), *TNFRSF11A* (Fw: ATCTGGGACGGTGCTGTAAC; Rv: GGCCTTGCCTGTATCACAAA), *TNFSF11* (Fw: TGATTCATGTAGGAGAATTAAACAGG; Rv: GATGTGCTGTGATCCAACGA), *BIRC3* (Fw: GGTAACAGTGATGATGTCAAATG; Rv: TAACTGGCTTGAACTTGACG), *ICAM1* (Fw: AACTGACACCTTTGTTAGCCACCTC; Rv: CCCAGTGAAATGCAAACAGGAC), *CCL2* (Fw: AGGTGACTGGGCATTGAT; Rv: GCCTCCAGCATGAAAGTCT), *IL8* (Fw: CTGCGCCAACACAGAAATTA; Rv: CATCTGGCAACCCTACAACA), *RELB* (Fw: CCCGACCTCTCCTCACTCTC; Rv: CAGGGTGACCGTGCTCAG), *NF-kB2* (Fw: GGCGGGCGTCTAAAATTCTG; Rv: TCCAGACCTGGGTTGTAGCA).

### Western Blot

Five hundred thousand cells extracted from PDX tumors were seeded in growth medium (5% FBS, EGF 10 ng/ml, hydrocortisone 0.5 μg/ml, insulin 5 μg/ml, cholera toxin 100 ng/ml, and penicillin/streptomycin) o/n and then changed to starving medium (0.5% FBS, EGF 10 ng/ml, hydrocortisone 0.5 μg/ml, insulin 5 μg/ml, cholera toxin 100 ng/ml, and penicillin/streptomycin) during 24 hours before RANKL stimulation (500 ng/ml; Amgen Inc). Extracts for immunoblots were prepared with modified RIPA buffer (50 mM Tris pH 7.4, 150 nM NaCl, 1% Triton NP-40, 0.25% sodium deoxycholate) containing PhosSTOP and Complete protease inhibitor cocktail (Roche). Forty µg of total protein determined with DC protein assay reagents (BIO-RAD) were mixed with loading buffer (final concentrations: 62 mM Tris pH 6.8, 12% glycerol, 2.5% SDS) and 5% β-mercaptoethanol. Proteins were resolved by SDS-PAGE and transferred to Immobilon-P 0.45 µm membranes (Millipore). Primary antibodies against P-p65 (Ser536, Cell Signaling), p65 (D14E12, Cell Signaling), P-IkBα (S32/36, Cell Signaling), IkBα (L35A5 Cell Signaling), and β-tubulin (ab21058, Abcam) were used. Blots were incubated with HRP-conjugated secondary antibodies (DAKO) and developed with ECL detection kit (Amersham Biosciences).

### RNA sequencing

Total RNA samples were processed with the “QuantSeq 3’ mRNA-Seq Library Prep Kit (FWD) for Illumina” (Lexogen, Cat.No. 015) with RNA Quality scores of 7.7 on average (range 4.2-9.2). Library generation was initiated by reverse transcription with oligodT priming, and a second strand synthesis was performed from random primers. Libraries were completed by PCR. cDNA libraries were purified, applied to an Illumina flow cell for cluster generation and sequenced on an Illumina instrument. Read adapters and poly A tails were removed with BBDuk v38.38. Then, human reads were separated from mice ones using Xenome v1.0.1 (37) and those classified as “human”, “both” or “ambiguous” were selected. Processed reads were analyzed with the Nextpresso pipeline v1.9.2.5 (38). Sequencing quality was checked with FastQC v0.11.7 and FastQ Screen v0.13.0. Reads were aligned to the human reference genome (GRCh38) with TopHat v2.0.10 using Bowtie v1.0.0.0 and Samtools v0.1.19.0 (-library-type fr-secondstrand), allowing three mismatches and twenty multihits. Read counts were obtained with HTSeq-count v0.6.1 (--stranded=yes) using the human gene annotation from GENCODE (gencode.v34.GRCh38.Ensembl100). Differential expression was performed with DESeq2, using a 0.05 FDR. Genes were ranked according to the log2 Fold Change and GSEAPreranked v2.2.2 was used to perform gene set enrichment analysis for Hallmark, Biocarta, Reactome and KEGG v7.1 signatures, setting 1000 gene set permutations and a classic enrichment statistic. Only those signatures with significant enrichment levels (*FDR q-value < 0.25*) were considered.

## Statistical analysis

Statistical analysis in the TMA collections were performed with the support of the IDIBELL and Nottingham University Statistical Assessment Services. Associations between IHC scores and clinicopathologic parameters were evaluated using Pearson’s Chi-Square test or Fisher’s exact test. BCSS, DMFS and DFS were analyzed using the Kaplan–Meier function, Cox regression analyses and the log rank test. Data analyses of mouse experiments were performed using GraphPad Prism software version 8. Regression analysis of the growth curve mean for in vivo treatments was performed. Analysis of the differences between two conditions was performed with a two-tailed Student’s t-test.

Bubble matrix plots were drawn using R (v4.0.3) and ggplot2 (v3.3.3). These plots represent the NES and FDR values reported by GSEA for some selected pathways in all tested comparisons. The color scale represents the NES: red denotes a NES > 0 and blue a NES < 0. The more intense the color, the more extreme the NES. In addition, the size of the bubble is proportional to the -log10 of the FDR. Thus, the bigger the dot, the smaller the FDR. Gene sets were classified according to Pearson’s R coefficient generated by public gene set databases (KEGG, Biocarta, Reactome, and Hallmarks).

### Study approval

All human samples were obtained following institutional guidelines and the study received approval form the corresponding institutional Ethics Committee in accordance with the declaration of Helsinki. This work obtained ethics approval to use the human tissue samples by the corresponding institutional review boards: Greater Manchester Central Research Ethics Committee reference number 15/NW/0685 (Nottingham); Hospital Universitario 12 de Octubre, number 11/137 (CNIO) and Hospital Universitario de Bellvitge, PR166/11071/015. Informed consent was obtained from all individuals prior to surgery to use their tissue materials in research. Written informed consent for PDX generation was obtained from all subjects.

All experimental animal procedures were performed according to Spanish regulations. All research involving animals was performed at the IDIBELL animal facility in compliance with protocols approved by the IDIBELL Committee on Animal Care and following national and European Union regulations.

## Discussion

BC is a complex disease resulting from genetic and environmental factors with high intra- and inter-tumor heterogeneity that varies from good to very poor prognosis (39), (40). The current biomarkers that nowadays still dictate BC prognosis and treatment include hormone receptors, HER2 and Ki67. Gene expression analysis provides the oncologists with a superior tool to guide treatments based on molecular subtypes (41). However, great heterogeneity persists in all the subtypes, in particular in the TNBC and “basal-like” tumors that translates into wide-range of responses to current and still limited treatments. Although the development of new therapies has improved the prognosis and survival rates of BC patients (42, 43), many tumors remain therapy-resistant or acquire resistance with time (44). In order to provide an efficient treatment, the search for new prognostic and predictive factors has become an essential task for the individualization of BC therapy (45).

In the present work, we aimed to evaluate the potential value of RANK and RANKL as clinical predictors of BC prognosis. Our analyses of RANK and RANKL expression in more than 1500 BC samples from four independent cohorts corroborated previous findings, with RANK expression being associated with ER^-^ tumors and RANKL rarely found in tumor cells (7, 8). Importantly, the large number of samples analyzed in our study allowed to define RANK expression as an independent poor prognosis factor in ER^-^ BC disease and postmenopausal patients, and able to predict worse BCSS in the whole BC population.

The distinct biology associated with RANK signaling according to ER status may explain why RANK predicts poor prognosis in ER^-^, but not in ER^+^ BC. RANK protein expression in the tumor cells of ER^+^ tumors was negatively associated with several proliferation/ DNA repair pathways; this counterintuitive finding is in line with our recent results showing that RANK overexpression induces senescence in the luminal mammary epithelial cells and associates with senescence in luminal BC from the TCGA, but not in basal-like tumors (46). However, it is important to acknowledge that RANK is expressed at lower levels and less frequently in ER^+^ compared to ER^-^ disease, which, together with the limitations of the RANK IHC, would underestimate the number of ER^+^ tumors with active RANK signaling. Moreover, the frequency of RANK positivity in the *NPS* collection was lower than in the more recent cohorts analyzed. Additional ER^+^ collections need to be evaluated, to rule out RANK expression in tumor cells as a prognosis factor of ER^+^ BC.

GSEA results in the ER^-^ tumors and RANKL/RANK-Fc treated-ER^-^ PDX tumors evidence a pleiotropic role of RANK signaling in BC, regulating multiple biological processes and oncogenic/inflammatory pathways together with tumor cell proliferation/differentiation, metabolism, immunity, and adhesion, in line with previous findings (10,17,31,47). It is striking that even in a severe immunodeficient environment such as that of the NSG mice, several immune-related pathways are regulated. This is in agreement with our recent findings, demonstrating that RANK expression in tumor cells is a key regulator of the tumor immune response in preclinical models, but also in the BC patients (31). RANK loss in tumor cells led to greater anti-tumor effects in immunocompetent compared to immunodeficient models. Thus, we are underestimating the therapeutic benefit of RANKL inhibitors when using PDX models. The functional studies in ER^-^ BC PDX support that RANK-RANKL signaling promotes tumor progression and recurrence of ER^-^ tumors, increasing tumor cell proliferation and stemness. However, given the immunomodulatory role for RANK signaling, it is likely that these effects will be greater when adding the denosumab-driven anti-tumor immune response. Importantly, despite BC heterogeneity and the diversity of the PDX models used, the transcriptomic analyses revealed a wide range of overlap in RANK-driven mechanisms, and, even more important, overlap with associations found in clinical samples and genes regulated by denosumab in BC patients. The main advantage of using PDX models is that they allow us to identify “biomarkers of response to denosumab in human BC”, without the confounding effects of infiltrating cells. These findings could prove useful for the selection of BC patients who may benefit from denosumab and the evaluation of drug responsiveness during treatment. Although it has been demonstrated that RANK^+^ BC show a higher response to chemotherapy (8), this is due to the increased frequency of RANK in ER^-^ tumors, which are the most responsive to chemotherapy (41). Our results suggest that RANK expression in ER^-^ tumors and TNBC is associated with worse response to chemotherapy, in particular, to the chemotherapy regimens that include taxanes. An alternative explanation is that RANK expression in ER^-^ tumors identifies patients with the worst outcome, irrespectively of the treatment regimen. Functional results in the PDX models, combining taxanes with RANK pathway inhibitors demonstrate enhanced benefit of the combination compared to chemotherapy alone, with lower rates of recurrence. However, results from the GeparX clinical trial demonstrated that the use of neoadjuvant denosumab in combination with Nab-Paclitaxel did not increase the pathological complete response in patients with early BC, not even in patients with RANK^+^ early BC, but survival remains to be evaluated (48).

Paradoxically to the well characterize role of RANK signaling as a mediator of progesterone in the healthy breast or preneoplasic lesions (10, 15), our results clearly demonstrate that RANK is a prognosis factor in ER^-^ postmenopausal BC, but not in premenopausal. It is well known that the drop in estrogen levels leads to a reduction in OPG and enhances activation of RANK signaling in the bone leading to osteoporosis (22). Although the role of OPG in the breast remains poorly explored, following the same rationale, one might expect enhanced activation of RANK signaling also in the breast tumors after menopause, particularly in ER^-^ tumors, which have the highest levels of RANK. Indeed, GSEA analyses revealed that multiple pathways related to TNF/NFκB, including RANKL pathway, were positively associated with RANK protein expression in postmenopausal, but not in premenopausal patients, and these pathways overlapped with those regulated in PDX after RANK modulation. If this hypothesis is confirmed, denosumab may show the highest therapeutic benefit in postmenopausal women with ER^-^ RANK^+^ breast tumors. The meta-analysis by the Early Breast Cancer Clinical Trialists’ Collaborative Group supports the idea that adjuvant treatment of early BC might be more efficacious with the addition of a bone-modifying agent, particularly in postmenopausal women or in combination with ovarian function suppression (49, 50).

Previous results from ABCSG18 revealed that adjuvant denosumab reduced the risk of clinical fractures and improved DFS of hormone receptor-positive postmenopausal BC patients receiving aromatase inhibitors (19), but this was not validated in the D-CARE trial (20). However, in these trials RANK expression or RANK pathway activation was not considered. Subsequent analyses categorizing the groups based on RANK expression and/or RANK pathway activation based on gene expression signatures such as the RANK metagene (31) or pathways identified in this study, is critical to fully comprehend the therapeutic potential of RANK pathway inhibitors in BC. Until June 2021, denosumab has been included in 248 clinical trials, 122 of them have been completed. From the finished studies, 10% was aimed at analyzing the effectiveness of denosumab in BC (51). However, most of these clinical trials did not consider RANK expression or activation as selection criteria. Our results point out the importance of conducting a retrospective study considering RANK pathway activation when evaluating outcome after denosumab.

In summary, in this work we have shown that RANK can be used as an independent marker of poor prognosis for postmenopausal patients with ER^-^ BC and poor response to chemotherapy and that activation of RANK signaling functionally contributes to tumor progression and aggressiveness in ER^-^ tumors, supporting the therapeutic potential of RANK pathway inhibitors in ER^-^ BC.

## Acknowledgements

We thank all the patients who contributed to this study, Robert Clarke, Bruno Simoes (University of Manchester), Joaquin Arribas (VHIO), Alana Welm (Huntsman Cancer institute), Violeta Serra (VHIO) and Carlos Caldas (CRUK, University of Cambridge) for providing PDX tumor samples for the analyses of RANK and RANKL expression. We thank Amgen for providing the N1H8 and M366 antibodies and the recombinant RANKL and RANK-Fc proteins. We thank the IDIBELL Animal Facility for their assistance with mouse colonies, Dr Esther Castaño, Beatriz Barroso and the scientific services of the University of Barcelona for their assistance with flow cytometry analyses, Fran Cimas, Elayne Hondares, Idoia Morilla for the analyses of TMA and PDX cohorts Manuel Gris for NSG maintenance. We thank Miguel Angel Pujana, Christian Tebe and Judith Peñafiel for the analyses of the *IDB* and *CNIO* cohorts.

## Author contributions

MC: Collection and/or assembly of data, data analysis and interpretation, manuscript writing, coordination. EMT: Collection and/or assembly of data, data analysis and interpretation, manuscript writing. AV, HPM, ASM, MJJS, JGM, MJ, AP, MTSM, MA, MT, ARG, SM, MQ, FAL-S, AMA, AS, ER: Collection and/or assembly of data, data analysis and interpretation. GPC: data analysis and interpretation, manuscript writing. EG-S: Conception and design, financial support, collection and/or assembly of data, data analysis and interpretation, manuscript writing, coordination. LED, MTL, AB: provide reagents (PDX and TMA slides).

## References

1. Bray F, et al. Global cancer statistics 2018: GLOBOCAN estimates of incidence and mortality worldwide for 36 cancers in 185 countries. CA Cancer J Clin. 2018;68(6):394–424.

2. Perou CM et al. Molecular portraits of human breast tumours. Nature. 2000;406(6797):747–52.

3. Cheang MCU et al. Ki67 index, HER2 status, and prognosis of patients with luminal B breast cancer. J Natl Cancer Inst. 2009;101(10):736–50.

4. Dent R et al. Triple-negative breast cancer: Clinical features and patterns of recurrence. Clin Cancer Res. 2007;13(15):4429–34.

5. Gluz O, et al. Triple-negative breast cancer - Current status and future directions. Vol. 20, Annals of Oncology. Oxford University Press; 2009. p. 1913–27.

6. Bray F, et al. Global cancer statistics 2018: GLOBOCAN estimates of incidence and mortality worldwide for 36 cancers in 185 countries. CA Cancer J Clin. 2018;68(6):394–424.

7. Palafox M et al. RANK induces epithelial-mesenchymal transition and stemness in human mammary epithelial cells and promotes tumorigenesis and metastasis. Cancer Res. 2012;72(11):2879–88.

8. Pfitzner BM et al. RANK expression as a prognostic and predictive marker in breast cancer. Breast Cancer Res Treat. 2014 Apr 16;145(2):307–15.

9. Azim HA et al. RANK-ligand (RANKL) expression in young breast cancer patients and during pregnancy. Breast Cancer Res. 2015;17(1):24.

10. Gonzalez-Suarez E et al. RANK ligand mediates progestin-induced mammary epithelial proliferation and carcinogenesis. Nature 2010;468(7320):103–7.

11. Fata JE et al. The osteoclast differentiation factor osteoprotegerin-ligand is essential for mammary gland development. Cell. 2000;103(1):41–50.

12. Beleut M et al. Two distinct mechanisms underlie progesterone-induced proliferation in the mammary gland. Proc Natl Acad Sci U S A. 2010;107(7):2989–94.

13. Wang J et al. Comment on “Progesterone/RANKL is a major regulatory axis in the human breast.” Sci Transl Med. 2013;5(215):215le4.

14. Joshi PA et al. Progesterone induces adult mammary stem cell expansion. Nature. 2010;465(7299):803–7.

15. Schramek D et al. Osteoclast differentiation factor RANKL controls development of progestin-driven mammary cancer. Nature. 2010;468(7320):98–102.

16. Nolan E et al. RANK ligand as a potential target for breast cancer prevention in BRCA1-mutation carriers. Nat Med. 2016;22(8):933–9.

17. Yoldi G et al. RANK signaling blockade reduces breast cancer recurrence by inducing tumor cell differentiation. Cancer Res. 2016;76(19):5857–69.

18. Miyazaki T, et al. A review of denosumab for the treatment of osteoporosis. Vol. 8, Patient Preference and Adherence. DOVE Medical Press Ltd.; 2014. p. 463–71.

19. Gnant M et al. Adjuvant denosumab in early breast cancer: Disease-free survival analysis of 3,425 postmenopausal patients in the ABCSG-18 trial. J Clin Oncol. 2018 May 20;36(15_suppl):500–500.

20. Coleman R et al. Adjuvant denosumab in early breast cancer (D-CARE): an international, multicentre, randomised, controlled, phase 3 trial. Lancet Oncol. 2020;21(1):60–72.

21. Curtis C et al. Dunning MJ, et al. The genomic and transcriptomic architecture of 2,000 breast tumours reveals novel subgroups. Nature. 2012;486(7403):346–52.

22. Streicher C et al. Estrogen Regulates Bone Turnover by Targeting RANKL Expression in Bone Lining Cells. Sci Rep. 2017;7(1).

23. Bruna A et al. A Biobank of Breast Cancer Explants with Preserved Intra-tumor Heterogeneity to Screen Anticancer Compounds. Cell. 2016;167(1):260–274.e22.

24. Derose YS et al. Tumor grafts derived from women with breast cancer authentically reflect tumor pathology, growth, metastasis and disease outcomes. Nat Med. 2011;17(11):1514–20.

25. Eyre R et al. Patient-derived mammosphere and xenograft tumour initiation correlates with progression to metastasis. J Mammary Gland Biol Neoplasia. 2016;21(3–4):99–109.

26. Zhang X et al. Establishment of Patient-Derived Xenograft (PDX) Models of Human Breast Cancer. Curr Protoc Mouse Biol. 2013;3(1):21–9.

27. Gómez-Miragaya J et al. Resistance to Taxanes in Triple-Negative Breast Cancer Associates with the Dynamics of a CD49f+ Tumor-Initiating Population. Stem Cell Reports. 2017;8(5):1392–407.

28. Gris-Oliver A et al. Genetic Alterations in the PI3K/AKT Pathway and Baseline AKT Activity Define AKT Inhibitor Sensitivity in Breast Cancer Patient-derived Xenografts. Clin Cancer Res. 2020 15;26(14):3720–31.

29. Santini D et al. Receptor activator of NF-kB (rank) expression in primary tumors associates with bone metastasis occurrence in breast cancer patients. PLoS One. 2011;6(4).

30. Gomez-Miragaya J et al. Tumor-initiating CD49f cells are a hallmark of chemoresistant triple negative breast cancer. Mol Cell Oncol. 2017;4(4):e1338208–e1338208.

31. Gómez-Aleza C et al. Inhibition of RANK signaling in breast cancer induces an anti-tumor immune response orchestrated by CD8+ T cells. Nat Commun. 2020;11(1):1–18.

32. Rakha EA et al. Prognostic significance of nottingham histologic grade in invasive breast carcinoma. J Clin Oncol. 2008;26(19):3153–8.

33. Elston Cw et al. Pathological prognostic factors in breast cancer. I. The value of histological grade in breast cancer: experience from a large study with long-term follow-up. Histopathology. 1991;19(5):403–10.

34. Abd El-Rehim DM et al. High-throughput protein expression analysis using tissue microarray technology of a large well-characterised series identifies biologically distinct classes of breast cancer confirming recent cDNA expression analyses. Int J Cancer. 2005;116(3):340–50.

35. Smalley MJ. Chapter 11 Isolation, Culture and Analysis of Mouse Mammary Epithelial Cells. Methods Mol Biol. 2010; 633:139–70.

36. Schindelin J, et al. Fiji: An open-source platform for biological-image analysis. Vol. 9, Nature Methods. Nat Methods; 2012. p. 676–82.

37. Conway T et al. Xenome-a tool for classifying reads from xenograft samples. Bioinformatics. 2012;28(12):172–8.

38. Graña O et al. Nextpresso: Next Generation Sequencing Expression Analysis Pipeline. Curr Bioinform. 2018;13(6):583–91.

39. Skol AD et al. The genetics of breast cancer risk in the post-genome era: Thoughts on study design to move past BRCA and towards clinical relevance. Vol. 18, Breast Cancer Research. BioMed Central Ltd.; 2016.

40. Turashvili G et al. Tumor heterogeneity in breast cancer. Vol. 4, Frontiers in Medicine. Frontiers Media S.A.; 2017 p. 227.

41. Yersal O et al. Biological subtypes of breast cancer: Prognostic and therapeutic implications. Vol. 5, World Journal of Clinical Oncology. Baishideng Publishing Group Co., Limited; 2014. p. 412–24.

42. Lin SX et al. Molecular therapy of breast cancer: Progress and future directions. Vol. 6, Nature Reviews Endocrinology. 2010. p. 485–93.

43. Schütz F, et al. Update Breast Cancer 2019 Part 4 - Diagnostic and Therapeutic Challenges of New, Personalised Therapies for Patients with Early Breast Cancer. Geburtshilfe Frauenheilkd. 2019;79(10):1079–89.

44. Gonzalez-Angulo AM et al. Overview of resistance to systemic therapy in patients with breast cancer. Vol. 608, Advances in Experimental Medicine and Biology. Springer, New York, NY; 2007 p. 1–22.

45. Weigel MT et al. Current and emerging biomarkers in breast cancer: Prognosis and prediction. Vol. 17, Endocrine-Related Cancer. Society for Endocrinology; 2010. p. R245–62.

46. Sandra Benítez A et al. RANK links senescence to stemness in the mammary epithelia, delaying tumor onset but increasing tumor aggressiveness. Dev Cell. 2021;56:1727–1741.e7.

47. Rao S, Sigl V et al. RANK rewires energy homeostasis in lung cancer cells and drives primary lung cancer. Genes Dev. 2017 Oct 15;31(20):2099–111.

48. GeparX: Denosumab (Dmab) as add-on to different regimen of nab-paclitaxel (nP)-anthracycline based neoadjuvant chemotherapy (NACT) in early breast… | OncologyPRO.

49. T C, Y L, A F. Antiresorptive agents’ bone-protective and adjuvant effects in postmenopausal women with early breast cancer. Br J Clin Pharmacol. 2019;85(6):1125–35.

50. Perrone F et al. Denosumab in early breast cancer: negative data and a call to action. Vol. 21, The Lancet Oncology. Lancet Publishing Group; 2020. p. 5–6.

51. Denosumab | Completed Studies - List Results - ClinicalTrials.gov.

