## Supplemental Figures for "RANK is an independent biomarker of poor prognosis in estrogen receptor-negative breast cancer and a therapeutic target in patient-derived xenografts"

### Figure S1

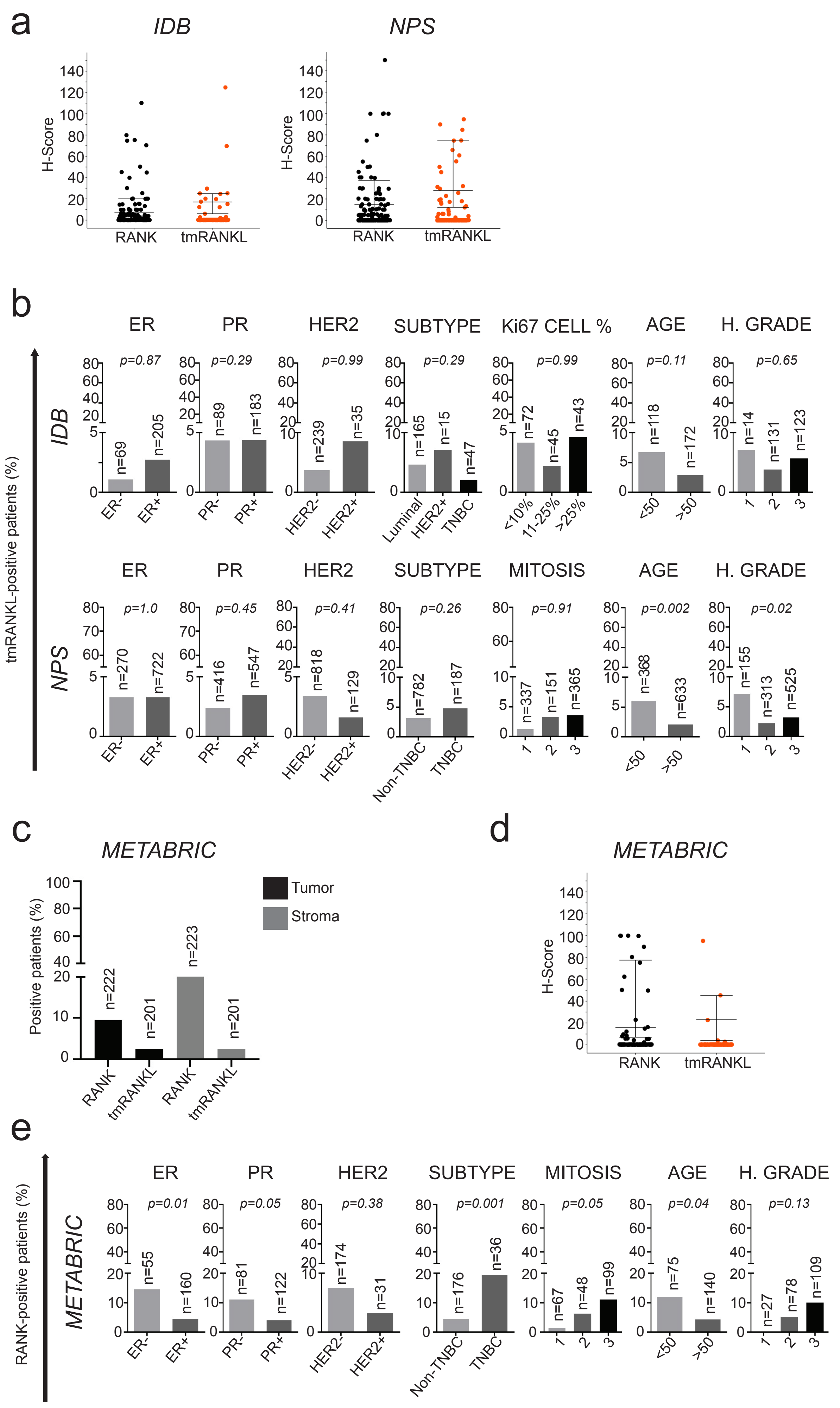
a
IDB
NPS
H-Score
RANK
tmRANKL
b
ER
PR
HER2
SUBTYPE
Ki67 CELL %
AGE
H. GRADE
IDB
NPS
tmRANKL-positive patients (%)
p=0.87
p=0.29
p=0.99
p=0.29
p=0.99
p=0.11
p=0.65
n=69
n=205
n=89
n=183
n=239
n=35
n=165
n=15
n=47
n=72
n=45
n=43
n=118
n=172
n=14
n=131
n=123
ER-
ER+
PR-
PR+
HER2-
HER2+
Luminal
TNBC
<10%
11-25%
>25%
<50
>50
MITOSIS
p=1.0
p=0.45
p=0.41
p=0.26
p=0.91
p=0.002
p=0.02
n=270
n=722
n=416
n=547
n=818
n=129
n=782
n=187
n=337
n=151
n=365
n=368
n=633
n=155
n=313
n=525
Non-TNBC
c
METABRIC
Positive patients (%)
Tumor
Stroma
n=222
n=201
n=223
n=201
d
METABRIC
e
METABRIC
RANK-positive patients (%)
p=0.01
p=0.05
p=0.38
p=0.001
p=0.05
p=0.04
p=0.13
n=55
n=160
n=81
n=122
n=174
n=31
n=176
n=36
n=67
n=48
n=99
n=75
n=140
n=27
n=78
n=109

**Fig. S1. Tumor tmRANKL expression is not associated with clinicopathologic factors in human breast adenocarcinomas.**

**(a)** H-Score values of tumor RANK and tm-RANKL from IDB and NPS collections. **(b)** Percentage of BC patients with tumor $tmRANKL^{+}$ tumors according to the indicated clinicopathologic parameters in the *IDB* and *NPS* cohorts. The total number of patients analyzed per parameter and *p-values* (calculated using the Pearson´s Chi-Square test (Exact Sig. 2- Side)) are indicated. **(c)** Percentage of patients expressing RANK or tmRANKL ($H > 0$) in tumor and stromal cells in BC samples from the *METABRIC* collection. The total number of patients scored for RANK and tmRANKL protein expression is indicated. **(d)** H-Score values of tumor RANK and tmRANKL from the *METABRIC* dataset. **(e)** Percentage of tumor $RANK^{+}$ BC patients according to the indicated clinicopathologic parameters in the *METABRIC* cohort. The total number of patients analyzed per parameter and *p-values* (calculated using the Pearson´s Chi-Square test (Exact Sig. 2-Side)) are indicated.

### Figure S2

a

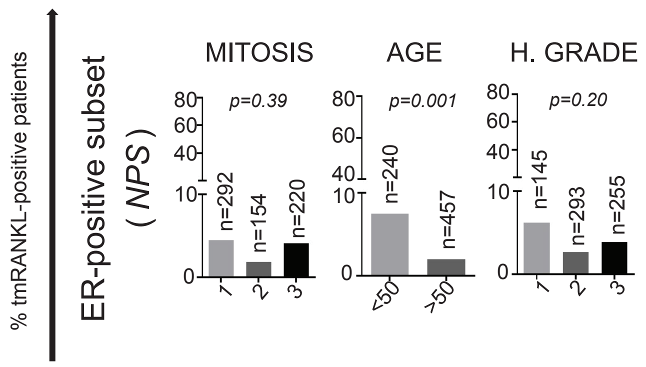
% tmRANKL-positive patients
ER-positive subset
( NPS )
MITOSIS
p=0.39
n=292
n=154
n=220
1
2
3
AGE
p=0.001
n=240
n=457
<50
>50
H. GRADE
p=0.20
n=145
n=293
n=255
1
2
3

b

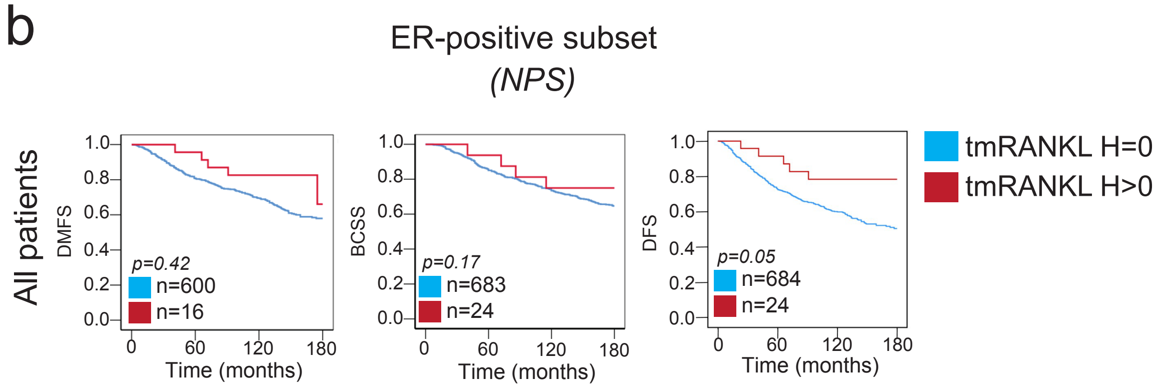
ER-positive subset
(NPS)
All patients
DMFS
p=0.42
n=600
n=16
Time (months)
BCSS
p=0.17
n=683
n=24
Time (months)
DFS
p=0.05
n=684
n=24
Time (months)
tmRANKL H=0
tmRANKL H>0

**Fig. S2. RANK tumor expression is not related to any clinicopathologic factor and RANKL associates with younger patients in ER± BC.**

**(a)** Percentage of tumor $RANK^+$ BC patients according to the indicated clinicopathologic parameters in the $ER^+$ subset from the *NPS*. The total number of patients analyzed per parameter and *p-values* (calculated using the Pearson´s Chi-Square tests (Exact Sig. 2-Side)) are indicated. **(b)** Percentage of tumor $tmRANKL^+$ BC patients according to the indicated clinicopathologic parameters in the $ER^+$ subset from the *NPS* collection. **(c)** DMFS, BCSS and DFS (15 years of follow-up) in the $ER^+$ subset from the *NPS* collection according to tmRANKL expression. **(a-b)** The total number of patients analyzed per parameter and *p-values* (calculated using the Pearson´s Chi-Square test (Exact Sig. 2-Side)) **(c)** and the Log-rank test (Mantel-Cox) are indicated.

### Figure S3

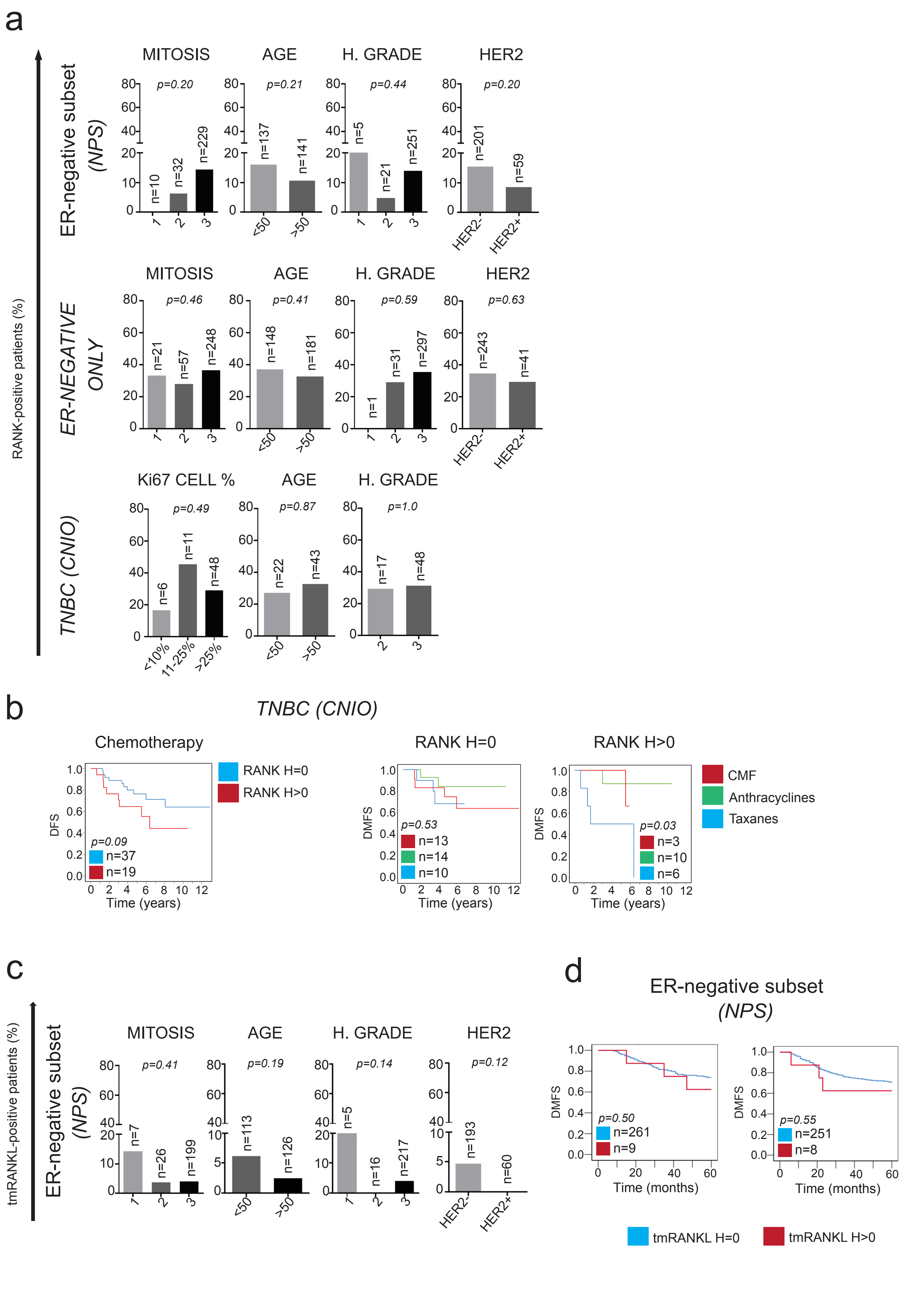
a
RANK-positive patients (%)
ER-negative subset (NPS)
MITOSIS p=0.20 n=10 n=32 n=229 1 2 3
AGE p=0.21 n=137 n=141 <50 >50
H. GRADE p=0.44 n=5 n=21 n=251 1 2 3
HER2 p=0.20 n=201 n=59 HER2- HER2+
ER-NEGATIVE ONLY
MITOSIS p=0.46 n=21 n=57 n=248 1 2 3
AGE p=0.41 n=148 n=181 <50 >50
H. GRADE p=0.59 n=1 n=31 n=297 1 2 3
HER2 p=0.63 n=243 n=41 HER2- HER2+
TNBC (CNIO)
Ki67 CELL % p=0.49 n=6 n=11 n=48 <10% 11-25% >25%
AGE p=0.87 n=22 n=43 <50 >50
H. GRADE p=1.0 n=17 n=48 2 3
b
TNBC (CNIO)
Chemotherapy
DFS
p=0.09 n=37 n=19
Time (years)
RANK H=0
RANK H>0
RANK H=0
DMFS
p=0.53 n=13 n=14 n=10
Time (years)
RANK H>0
DMFS
p=0.03 n=3 n=10 n=6
Time (years)
CMF
Anthracyclines
Taxanes
c
tmRANKL-positive patients (%)
ER-negative subset (NPS)
MITOSIS p=0.41 n=7 n=26 n=199 1 2 3
AGE p=0.19 n=113 n=126 <50 >50
H. GRADE p=0.14 n=5 n=16 n=217 1 2 3
HER2 p=0.12 n=193 n=60 HER2- HER2+
d
ER-negative subset (NPS)
DMFS
p=0.50 n=261 n=9
Time (months)
DMFS
p=0.55 n=251 n=8
Time (months)
tmRANKL H=0
tmRANKL H>0

**Fig. S3. RANK or RANKL tumor expression are not related to any clinicopathologic factor in ER- BC but RANK associates with poor response to chemotherapy including taxanes**

**(a)** Percentage of tumor RANK+ BC patients according to the indicated clinicopathologic parameters in the ER- subset from *NPS*, in the *ER-NEGATIVE ONLY* and *TNBC CNIO* collections. **(b)** DFS in the *TNBC CNIO* collection (12 years of follow-up) after chemotherapy according to RANK expression and DMFS in RANK- (H = 0) and RANK+ (H > 0) tumor samples from *TNBC CNIO* collection according to the chemotherapy regimen (12 years of follow-up): CMF treatment (cyclophosphamide, methotrexate, 5-fluorouracil), anthracycline treatment (FAC/FEC: 5-flurouracil, adriamycin, cyclophosphamide or 5-fluorouracil, epirubicin and cyclophosphamide) and taxane group (CMF/FAC/FEC plus taxanes). **(c)** Percentage of tumor tmRANKL+ BC patients according to the indicated clinicopathologic parameters in the *ER-negative subset* from *NPS*. **(d)** DFS and DMFS in the *ER- subset* from *NPS* (5 years of follow-up) according to the tmRANKL expression (tmRANKL- (H-Score=0) or tm-RANKL+ (H-Score>0)). **(a, c)** The total number of patients analyzed per parameter and *p-values* (calculated using the Pearson´s Chi-Square test (Exact Sig. 2-side)) are indicated. **(b, d)** The total number of patients analyzed per parameter and *p-values* (calculated using the Log-rank test (Mantel-Cox)) are indicated.

### Figure S4

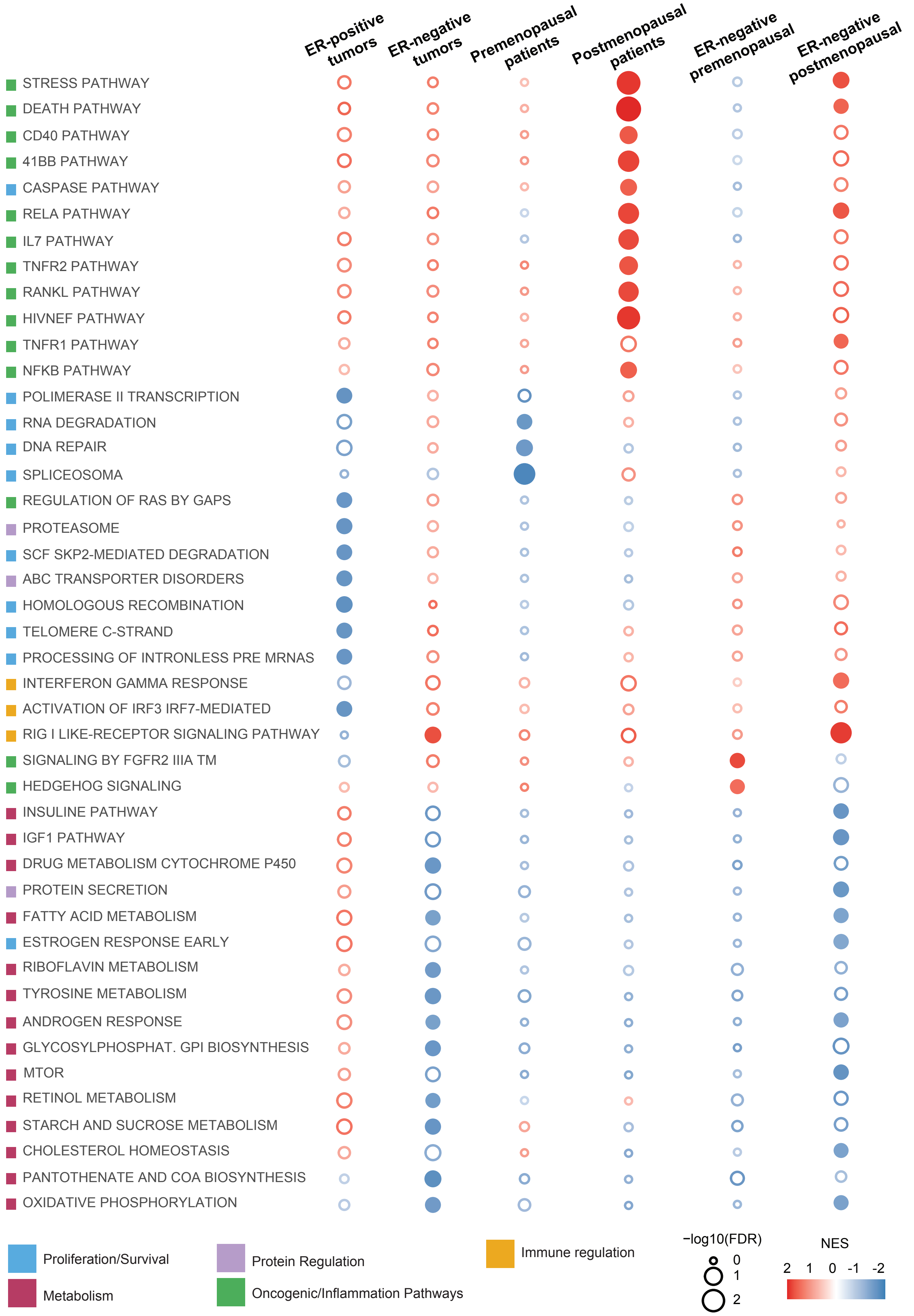

ER-positive tumors
ER-negative tumors
Premenopausal patients
Postmenopausal patients
ER-negative premenopausal
ER-negative postmenopausal
STRESS PATHWAY
DEATH PATHWAY
CD40 PATHWAY
41BB PATHWAY
CASPASE PATHWAY
RELA PATHWAY
IL7 PATHWAY
TNFR2 PATHWAY
RANKL PATHWAY
HIVNEF PATHWAY
TNFR1 PATHWAY
NFKB PATHWAY
POLIMERASE II TRANSCRIPTION
RNA DEGRADATION
DNA REPAIR
SPLICEOSOMA
REGULATION OF RAS BY GAPS
PROTEASOME
SCF SKP2-MEDIATED DEGRADATION
ABC TRANSPORTER DISORDERS
HOMOLOGOUS RECOMBINATION
TELOMERE C-STRAND
PROCESSING OF INTRONLESS PRE MRNAS
INTERFERON GAMMA RESPONSE
ACTIVATION OF IRF3 IRF7-MEDIATED
RIG I LIKE-RECEPTOR SIGNALING PATHWAY
SIGNALING BY FGFR2 IIIA TM
HEDGEHOG SIGNALING
INSULINE PATHWAY
IGF1 PATHWAY
DRUG METABOLISM CYTOCHROME P450
PROTEIN SECRETION
FATTY ACID METABOLISM
ESTROGEN RESPONSE EARLY
RIBOFLAVIN METABOLISM
TYROSINE METABOLISM
ANDROGEN RESPONSE
GLYCOSYLPHOSPHAT. GPI BIOSYNTHESIS
MTOR
RETINOL METABOLISM
STARCH AND SUCROSE METABOLISM
CHOLESTEROL HOMEOSTASIS
PANTOTHENATE AND COA BIOSYNTHESIS
OXIDATIVE PHOSPHORYLATION
Proliferation/Survival
Metabolism
Protein Regulation
Oncogenic/Inflammation Pathways
Immune regulation
−log10(FDR)
0
1
2
NES
2 1 0 -1 -2

**Fig. S4. Distinct biology of RANK signaling according to ER expression and menopausal status.**

Bubble matrix represents GSEA results of associated genes with RANK protein expression in the *METABRIC* collection classified by ER expression and menopausal status. Empty bubbles represent *FDR* < *0.25*. The matrix illustrates the NES and *FDR* values. Color legend indicate the main biological process associated.

### Figure S5

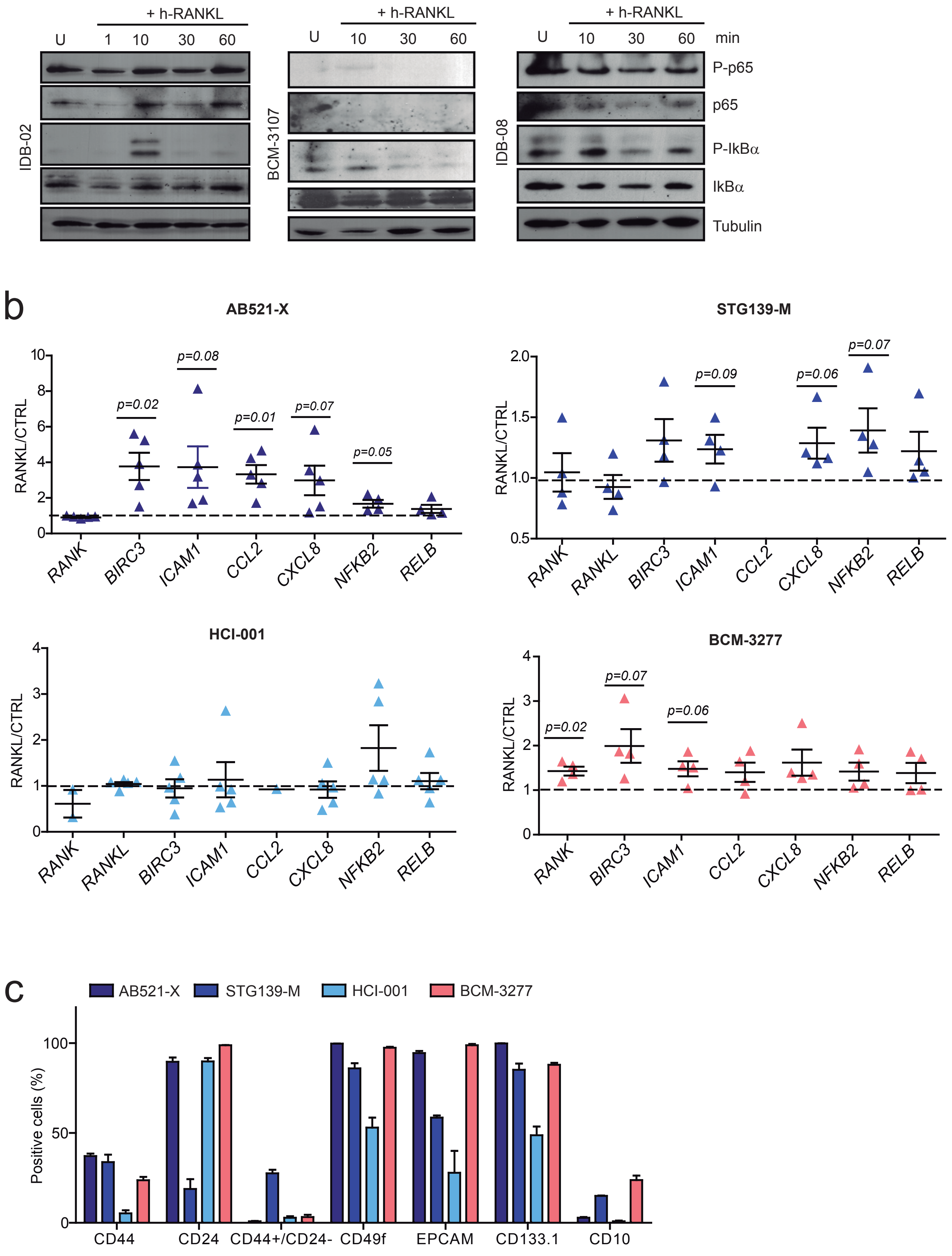

+ h-RANKL
U 1 10 30 60
IDB-02
U 10 30 60
BCM-3107
U 10 30 60 min
IDB-08
P-p65
p65
P-IkBα
IkBα
Tubulin
b
AB521-X
STG139-M
HCI-001
BCM-3277
RANKL/CTRL
p=0.02
p=0.08
p=0.01
p=0.07
p=0.05
p=0.09
p=0.06
p=0.07
p=0.02
p=0.07
p=0.06
RANK
RANKL
BIRC3
ICAM1
CCL2
CXCL8
NFKB2
RELB
c
AB521-X
STG139-M
HCI-001
BCM-3277
Positive cells (%)
CD44
CD24
CD44+/CD24-
CD49f
EPCAM
CD133.1
CD10

**Fig. S5. RANK signaling is active in a subset of BC PDXs.**

**(a)** Western blot analyses of P-p65, P-IKBα and corresponding total proteins after RANKL stimulation in the indicated PDXs. Tubulin was used as a loading control. **(b)** Gene expression analyses of the indicated NFκB target genes in PDX tumor organoids, 24 hours after RANKL stimulation. Expression levels relative to the untreated controls is shown. Each dot represents organoids from an independent BC PDX tumor. **(c)** Percentage of cells expressing the indicated surface markers for the four PDXs analyzed.

### Figure S6

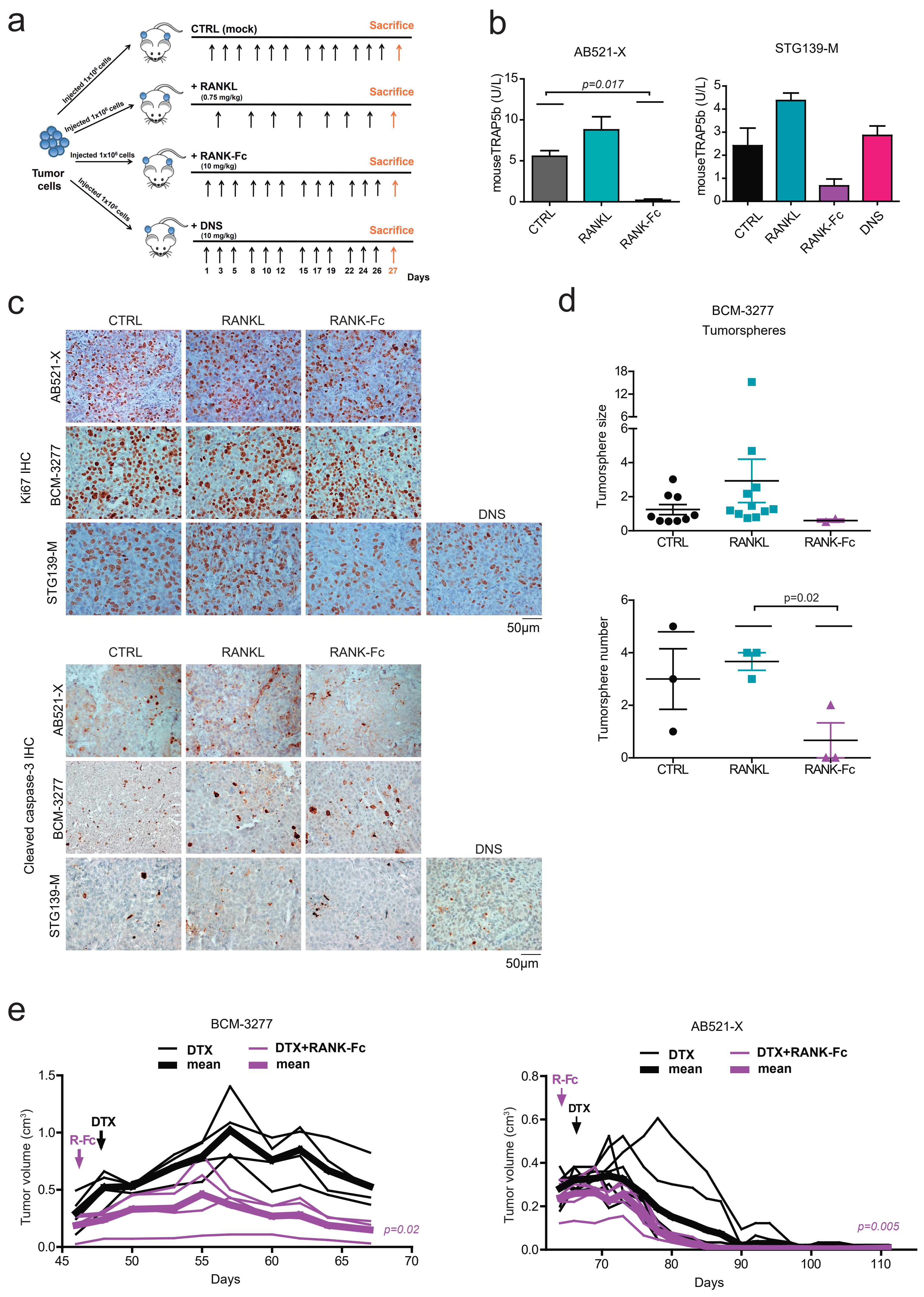
a
CTRL (mock)
Sacrifice
+ RANKL
(0.75 mg/kg)
+ RANK-Fc
(10 mg/kg)
+ DNS
(10 mg/kg)
Tumor cells
Injected 1x10⁶ cells
1 3 5 8 10 12 15 17 19 22 24 26 27
Days
b
AB521-X
STG139-M
mouseTRAP5b (U/L)
p=0.017
CTRL
RANKL
RANK-Fc
DNS
c
CTRL
RANKL
RANK-Fc
DNS
AB521-X
BCM-3277
STG139-M
Ki67 IHC
Cleaved caspase-3 IHC
50µm
d
BCM-3277
Tumorspheres
Tumorsphere size
Tumorsphere number
p=0.02
e
BCM-3277
AB521-X
DTX
mean
DTX+RANK-Fc
mean
R-Fc
Tumor volume (cm³)
Days
p=0.02
p=0.005

**Fig. S6. Modulation of RANK signaling in BC PDXs.**

**(a)** Schematic representation of *in vivo* treatments. Dissociated tumor cells mixed 1:1 with Matrigel Basement were transplanted orthotopically in NSG mice, when tumors reached 5 mm of diameter mice were randomized for mock (CTRL), h-RANKL (0.75 mg/kg, 4-6 doses, twice times per week), h-RANK-Fc treatment (10 mg/kg, three times per week), denosumab (DNS, 10 mg/kg, three times per week). Tumor development was monitored once per week. Tumor volume was calculated multiplying π x length x width$^2$/6 in cm. After 24 hours that the treatment was completed mice were sacrificed, and tumor cell proliferation and survival were measured and RNAseq performed. Trap5b levels were analyzed in serum. Tumor cells were isolated and tested for ALDH activity and tumorsphere forming ability. **(b)** Trap5b levels as determined by ELISA in mouse serum at the end of treatment. **(c)** Representative images of Ki67 and cleaved caspase-3 staining measured by IHC; see quantifications in Fig. 5b. **(d)** Size and total number of secondary tumorspheres stimulated *in vitro* with RANKL or RANK-Fc as indicated, stimulations were performed in triplicates and each dot represents a replicate. Three independent pictures per replicate were quantified for tumorsphere size. *P-value* was calculated using a two tailed t-student test **(e)** Tumor growth curves ((π x length x width$^2$)/6) of indicated PDXs after treatment with docetaxel (DTX, 20 mg/kg, once per week) in combination with RANK-Fc. Treatment started as indicated by the arrows. Each thin curve represents one single tumor and each thick curve represents the mean of all tumors implanted. Linear regression analysis was performed and two-tailed *p-value* is shown.

### Figure S7

a

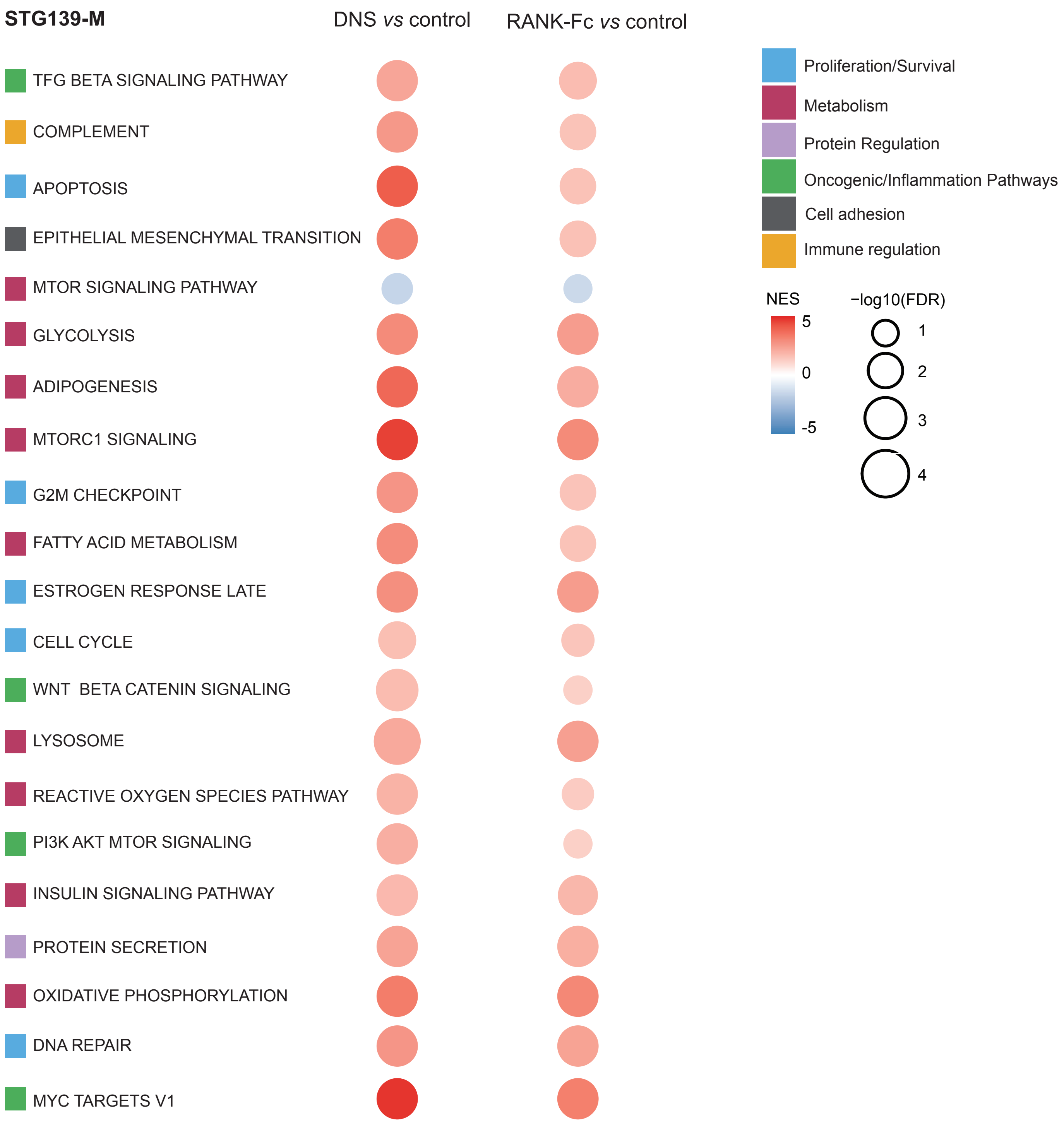

STG139-M
DNS vs control
RANK-Fc vs control
TFG BETA SIGNALING PATHWAY
COMPLEMENT
APOPTOSIS
EPITHELIAL MESENCHYMAL TRANSITION
MTOR SIGNALING PATHWAY
GLYCOLYSIS
ADIPOGENESIS
MTORC1 SIGNALING
G2M CHECKPOINT
FATTY ACID METABOLISM
ESTROGEN RESPONSE LATE
CELL CYCLE
WNT BETA CATENIN SIGNALING
LYSOSOME
REACTIVE OXYGEN SPECIES PATHWAY
PI3K AKT MTOR SIGNALING
INSULIN SIGNALING PATHWAY
PROTEIN SECRETION
OXIDATIVE PHOSPHORYLATION
DNA REPAIR
MYC TARGETS V1
Proliferation/Survival
Metabolism
Protein Regulation
Oncogenic/Inflammation Pathways
Cell adhesion
Immune regulation
NES
5
0
-5
−log10(FDR)
1
2
3
4


b

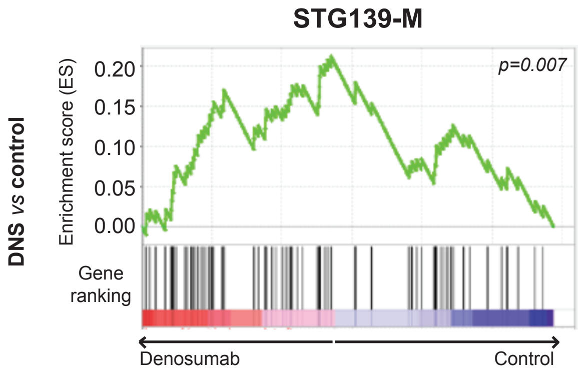

STG139-M
p=0.007
DNS vs control
Enrichment score (ES)
0.20
0.15
0.10
0.05
0.00
Gene ranking
Denosumab
Control

**Fig. S7. RANK inhibition regulates proliferation and oncogenic pathways mainly in BC PDXs.**

**(a)** Bubble matrix represents gene set enrichment analysis (GSEA) results of associated genes after *in vivo* treatment with denosumab (DNS) and RANK-Fc (RFc) in the STG139-M model. The matrix illustrates the NES and *FDR* values. The color scale represents the NES: red denotes a NES > 0 and blue a NES < 0. The size of the bubble is proportional to the -log10 of the *FDR*. Signatures belong to Hallmark, Biocarta, Reactome and KEGG collections. Colors **(b)** GSEA and genes upregulated by denosumab treatment in the D-BEYOND clinical trial (Gómez-Aleza et al. 2020) in STG139M treated with denosumab (DNS).
